# Overlapping and specialized roles of tomato phytoene synthase isoforms PSY1 and PSY2 in carotenoid and ABA production

**DOI:** 10.1101/2022.08.11.503628

**Authors:** Miguel Ezquerro, Esteban Burbano, Manuel Rodriguez-Concepcion

## Abstract

Carotenoids are plastidial isoprenoids required for photosynthesis and production of hormones such as abscisic acid (ABA) in all plants. In tomato (*Solanum lycopersicum*), carotenoids also provide color to flowers and ripe fruit. Phytoene synthase (PSY) catalyzes the first and main flux-controlling step of the carotenoid pathway. Three PSY isoforms are present in tomato, PSY1 to 3. Mutants have shown that PSY1 is the isoform providing carotenoids for fruit pigmentation but it is dispensable in photosynthetic tissues. No mutants are available for PSY2 or PSY3, but their expression profiles suggest a main role for PSY2 in leaves and PSY3 in roots. To further investigate isoform specialization with genetic tools, we created tomato edited lines defective in PSY1 and PSY2 in the MicroTom background. The albino phenotype of lines lacking both PSY1 and PSY2 confirmed that PSY3 does not contribute to carotenoid biosynthesis in shoot tissues. Our work further shows that carotenoid production in tomato shoots relies on both PSY1 and PSY2 but with different contributions in different tissues. PSY2 is the main isoform for carotenoid biosynthesis in leaf chloroplasts, but the supporting role of PSY1 is particularly important under high light. PSY2 also contributes to the production of carotenoids in flower petals and, to a lower extent, fruit chromoplasts. Most interestingly, our results demonstrate that fruit growth and ripening is controlled by ABA produced in the pericarp from PSY1-derived precursors whereas PSY2 provides precursors for ABA synthesis in seeds to control germination.

## Introduction

Carotenoids are a group of isoprenoid molecules synthetized by all photosynthetic organisms and some non-photosynthetic bacteria and fungi (Rodriguez-Concepcion et al., 2018; Sun et al., 2018). Carotenoids are essential micronutrients in our diet as precursors of retinoids such as vitamin A. Their characteristic colors in the range of yellow to orange and red also make them economically relevant as natural pigments in the chemical, pharma and agrofood industry. In plants, carotenoids are essential for photosynthesis (by contributing to the assembly of the photosynthetic apparatus and by participating in light harvesting) and for photoprotection (by dissipating the excess of light energy as heat and by scavenging free radicals). They also provide color to some non-photosynthetic tissues such as flower petals and ripe fruit to attract animals for pollination and seed dispersal. Besides, carotenoids are precursors of the phytohormones abscisic acid (ABA) and strigolactones (SL) and other biologically active signals involved in plastid-to-nucleus communication (e.g., beta-cyclocitral) and environmental interactions (e.g., apocarotenoids modulating root mycorrhization), among other processes (Moreno et al., 2021; Sun et al., 2018).

Carotenoids in plants are produced in plastids from geranylgeranyl diphosphate (GGPP) produced by the methylerythritol 4-phosphate (MEP) pathway (Fig. 1). GGPP is also used to produce other essential isoprenoids in the plastid, including plastoquinone, phylloquinone, tocopherols and chlorophylls (Rodriguez-Concepcion et al., 2018). The first committed step of carotenoid biosynthesis is the condensation of two GGPP molecules to produce phytoene (Fig. 1A). This step is catalyzed by phytoene synthase (PSY), the main flux-controlling enzyme of the carotenoid pathway (Cao et al., 2019; Zhou et al., 2022). Several desaturation and isomerization steps convert uncolored phytoene into red lycopene. From lycopene, carotenoid synthesis branches out depending on the type of cyclization of the ends of the lycopene carbon chain. The production of two β rings at the two ends of the chain produces β-carotene (β,β branch) while the production of one β ring and one ε ring produces α-carotene (β,ε branch). Oxygenation of the rings of carotenes produces xanthophylls such as violaxanthin and neoxanthin (β,β branch) or lutein (β,ε branch) (Fig. 1).

**Fig. 1.**
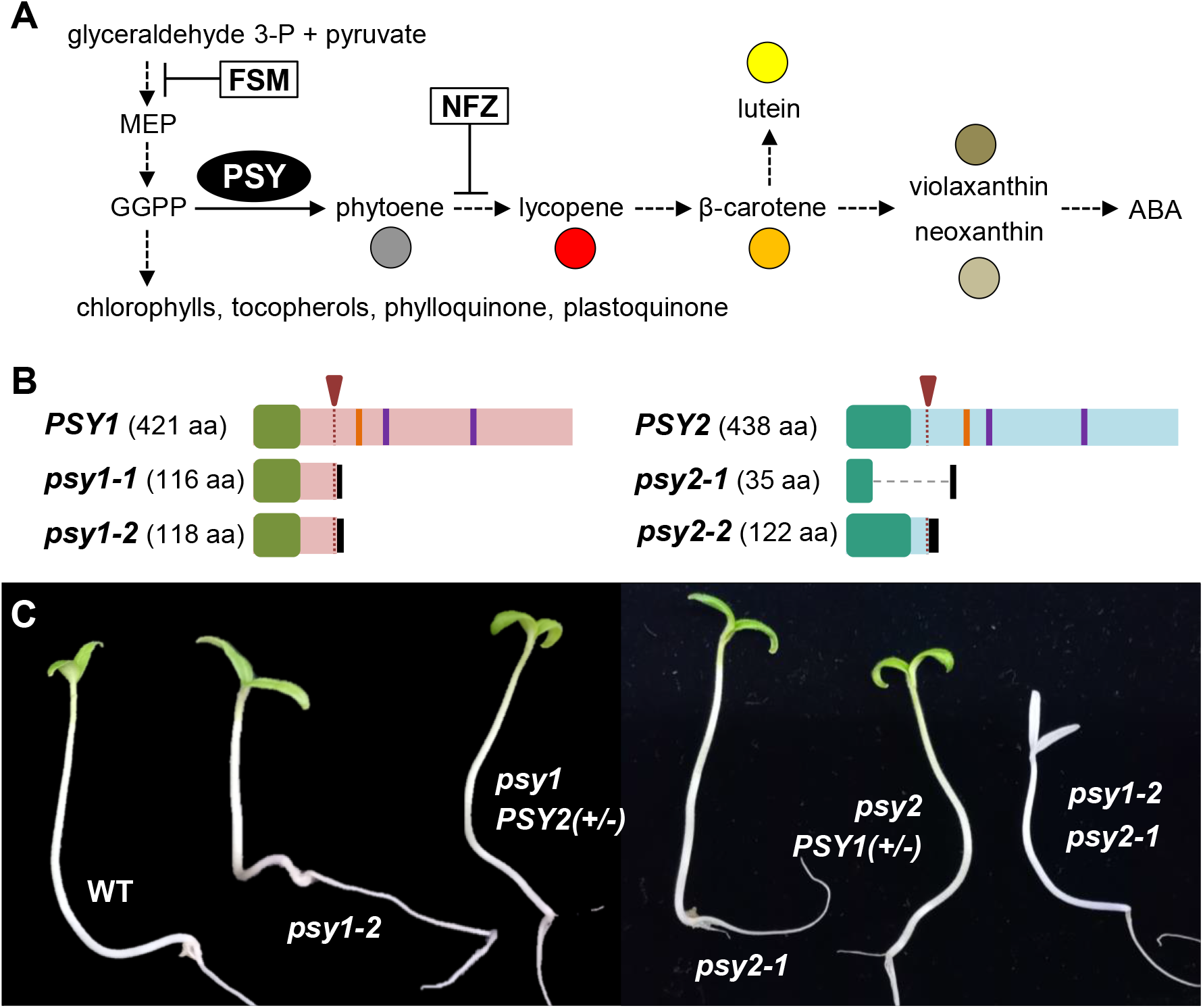
Carotenoid pathway and tomato mutants. **(A)** Carotenoid biosynthesis pathway. Dashed arrows represent multiple steps. The reaction catalyzed by phytoene synthase (PSY) is marked, and steps interrupted by inhibitors fosmidomycin (FSM) and norflurazon (NFZ) are indicated. Each individual carotenoid is represented by the indicated color (circle) in the corresponding plots representing their levels. **(B)** Scheme representing the wild-type PSY1 and PSY2 proteins and the mutant versions generated in the corresponding CRISPR-Cas9-generated alleles (see Fig. S1–S3 for further details). The region targeted by the designed sgRNAs is indicated with a red arrowhead and a dotted line. Orange and purple bars mark the position of conserved domains required for PSY activity (hydrofobic flap and Asp-rich domains, respectively). Green boxes represent plastid transit peptides. Black boxes represent the protein sequence resulting after a frame-shift in the mutants. The large deletion generated in the *psy2-1* allele is shown with a dashed line. **(C)** Representative seven-day-old seedlings of the indicated genotypes resulting from a cross of *psy1-2* and *psy2-1* mutants.

Tomato (*Solanum lycopersicum*) is a very well-suited model system to study carotenoid biosynthesis. Like all plants, tomato produces carotenoids for photosynthesis and photoprotection in chloroplasts and uses them as precursors to produce ABA and SLs in photosynthetic and non-photosynthetic tissues. But unlike Arabidopsis (*Arabidopsis thaliana*) and other plant models, tomato accumulates high levels of carotenoids in specialized plastids named chromoplasts, which are present in flower petals and ripe fruit. Also different from Arabidopsis, which only has a single PSY (At5g17230), the tomato genome harbors three PSY-encoding genes: *PSY1* (Solyc03g031860), *PSY2* (Solyc02g081330), and *PSY3* (Solyc01g005940) (Giorio et al., 2008; Stauder et al., 2018). While PSY1 and PSY2 are similar proteins that share conserved sequences and have a common origin (Cao et al., 2019; Giorio et al., 2008), PSY3 belongs to a different widespread clade restricted to dicots (Stauder et al., 2018). Tomato lines defective in PSY1 have been reported as *yellow-flesh* (*r*) mutants (Fray and Grierson 1993; Kachanovsky et al., 2012; Kang et al., 2014; Karniel et al., 2022), silenced lines (Bird et al., 1991; Bramley et al., 1992.; Fantini et al., 2013; Fraser et al., 1999) and CRISPR-Cas9-edited lines (D’Ambrosio et al., 2018), but lines impaired in PSY2 or PSY3 have not been described yet. Based on gene expression data and phenotypic features of PSY1-defective lines, it was proposed that PSY3 function might be restricted to roots whereas PSY1 and PSY2 differentially support carotenogenesis in shoot tissues: PSY1 for pigmentation in chromoplasts and PSY2 for photosynthesis in chloroplasts (Fraser et al., 1999; Giorio et al., 2008; Hirschberg 2001; Stauder et al., 2018). However, other sources of evidence suggest that isoform specialization is not complete. For example, the low but statistically significant upregulation of *PSY1* during seedling de-etiolation (when carotenoids are essential for the proper assembly of the photosynthetic apparatus and for photoprotection) and the high levels of *PSY2* transcripts in flower petals (where accumulation of xanthophylls is responsible for their characteristic yellow color) allows to hypothesize that both isoforms might participate in carotenoid biosynthesis in chloroplasts and chromoplasts (Barja et al., 2021; Giorio et al., 2008). To genetically test this hypothesis, we created tomato edited lines defective in PSY1 and PSY2 in the same tomato background (MicroTom, a widely used accession in molecular biology labs all over the world) and compared their physiological and metabolic phenotypes. The albino phenotype of lines defective in both PSY1 and PSY2 confirmed that PSY3 does not contribute to carotenoid biosynthesis in shoot tissues. Our work further confirmed that PSY2 is the main isoform supporting chloroplast carotenoid biosynthesis but uncovered a supporting role for PSY1 under conditions requiring an extra supply of carotenoids such as high light exposure. PSY1 was confirmed to be the main isoform in charge of phytoene production for carotenoid pigments in the chromoplasts of flower petals and fruit pericarp. Most interestingly, lower carotenoid levels resulted in a preferential reduction of ABA levels in the fruit pericarp but not in the seeds of the *psy1* mutant, whereas loss of PSY2 caused a major reduction of ABA in seeds. This differential ABA decrease in *psy1* and *psy2* mutants allowed to establish a specific contribution of pericarp ABA to fruit growth and ripening and seed ABA to seed germination.

## Results

### Loss of both PSY1 and PSY2 causes an albino-lethal phenotype

To generate plants defective in PSY1 or/and PSY2 in the MicroTom background, we designed one single guide RNA (sgRNA) annealing on the start of the first translated exon for each gene using the online tool CRISPR-P 2.0 (Liu et al., 2017). Two independent alleles with premature translation stop codons were selected for each gene and named *psy1-1, psy1-2, psy2-1* and *psy2-2* (Fig. 1B and Fig. S1-S3). For subsequent experiments we selected homozygous lines of each allele without the Cas9-encoding transgene. In the case of *psy1-1* and *psy1-2* alleles, we observed paler yellow flowers and pale orange fruits (Fig. S4), which are previously described phenotypes of *r* tomato lines, hence confirming that both were true PSY1-defective mutants. No distinctive phenotype was observed in the case of *psy2-1* and *psy2-2* lines. Analysis of transcript levels in fruits by RT-qPCR showed that loss of one of the isoforms did not influence the expression of the remaining genes (Fig. S4).

To assess the impact of simultaneous disruption of PSY1 and PSY2, we crossed lines defective in PSY1 (*psy1-2*, as female) and PSY2 (*psy2-1*, as male). Double heterozygous F_1_ plants with normal yellow flowers and red fruits were obtained and allowed to self-pollinate. Among the segregating F_2_ population we found several albino seedlings with a Mendelian proportion (1/16) consistent with this phenotype being the result of the loss of both PSY1 and PSY2 in double mutant individuals (Fig. 1C). The rest of the seedlings of the F_2_ population displayed a normal green phenotype indistinguishable from the MicroTom wild-type (WT). PCR-based genotyping of several individuals (Fig. S5) confirmed that green seedlings showed at least one WT copy of either *PSY1* or *PSY2* whereas all albino seedlings were double homozygous mutants. These results indicate that both PSY1 and PSY2 (but not PSY3) are essential for the production of carotenoids supporting seedling establishment and photosynthetic shoot development. Consistently, *PSY3* transcripts are hardly detectable in shoot tissues whereas *PSY1* and *PSY2* transcripts are abundant in all tissues of the tomato plant (Barja et al., 2021; Giorio et al., 2008; Stauder et al. 2018) (Fig. S6).

### PSY2 is supported by PSY1 to produce carotenoids for photoprotection in leaves

To test whether carotenoid levels were reduced in leaves of single *psy1* and *psy2* mutant lines, we collected young emerging leaves from plants grown for 18 days under long-day conditions in the greenhouse and used them for HPLC analysis of carotenoids and chlorophylls (Fig. 2). Despite WT and mutant plants were phenotypically identical (Fig. 2A), a slight reduction in carotenoid levels was detected in mutant leaves compared to WT controls (Fig. 2B). Chlorophylls were not as reduced as carotenoids (Fig. 2B). These results suggest that both PSY1 and PSY2 can produce carotenoids in chloroplasts under normal growth conditions, as the loss of one of the isoforms can be similarly rescued by the activity of the remaining isoform. Most interestingly, photosynthetic performance was only significantly reduced in *psy2* mutant alleles, as estimated from effective quantum yield of photosystem II (ΦPSII) measurements (Fig. 2C).

**Fig. 2.**
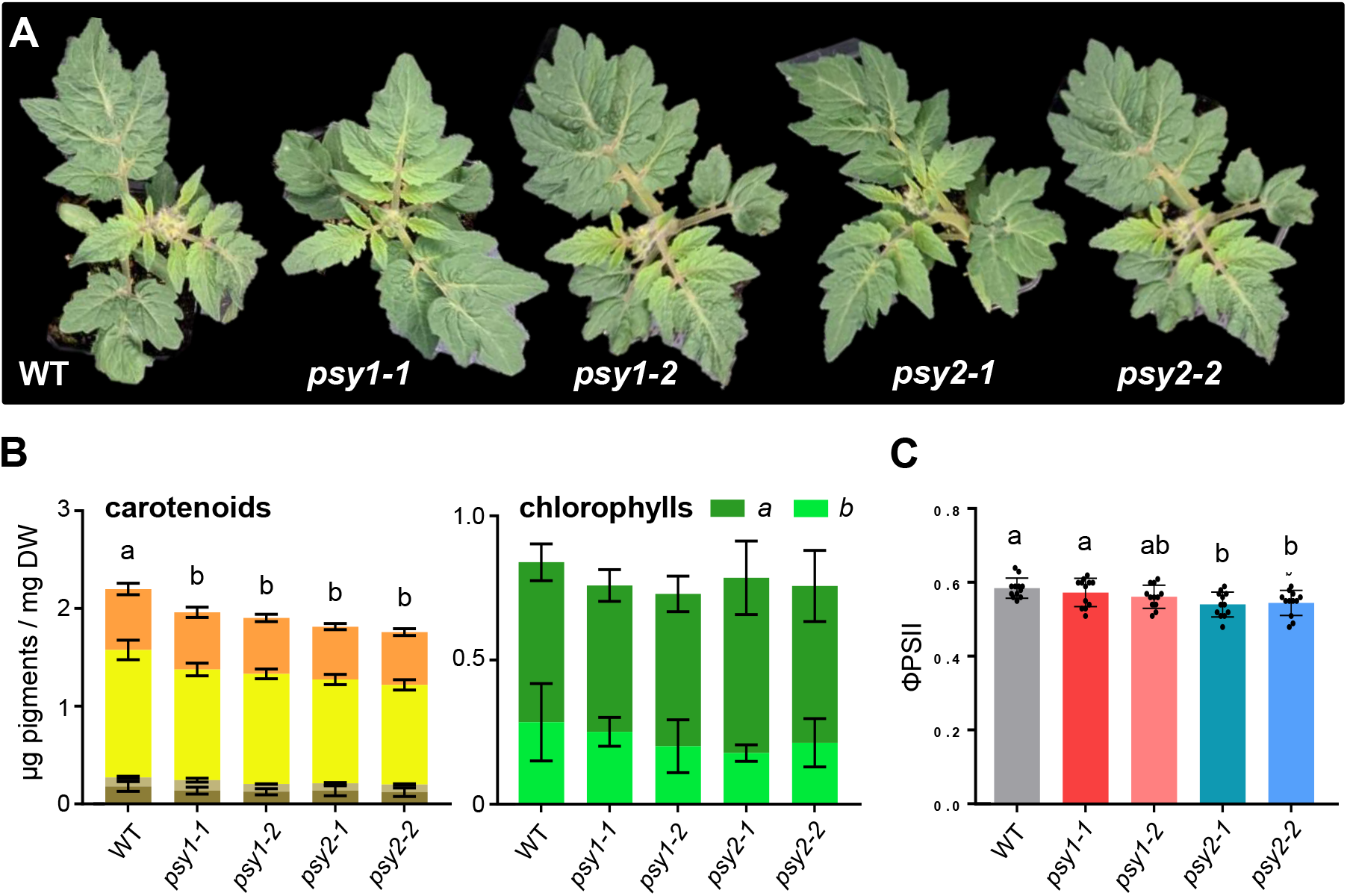
Tomato mutants defective in PSY1 or PSY2 show lower carotenoid levels under normal growth conditions. **(A)** Representative images of 4-week-old plants of the indicated lines. **(B)** Total levels of carotenoids and chlorophylls in young leaves of WT and mutant plants like those shown in (A). In the carotenoid plot, colors correspond to the species shown in Fig. 1A. Mean and SD of n≥3 independent biological replicates are shown. DW, dried weight. **(C)** Effective quantum yield of photosystem II (ɸPSII) in young leaves like those used in (B). Individual values (black dots) and well as mean and SD are shown, and they correspond to four different leaf areas from three different plants. In (B) and (C), bar letters represent statistically significant differences (*P* < 0.05) among means according to post hoc Tukey’s tests run when one way ANOVA detected different means.

The main role of carotenoids in photosynthetic organs such as leaves is photoprotection against photooxidative damage associated to intense light. In particular, carotenoids can dissipate the excess of light energy as heat through a process known as non-photochemical quenching (NPQ). Consistent with this essential function of leaf carotenoids, when 10-day-old tomato plants grown under normal light (NL) conditions (50 µmol photons m^−2^ s^−1^) were transferred to high light (HL) conditions (300 µmol photons m^−2^ s^−1^) for 5 days, expression of genes encoding PSY1 and PSY2 and concomitant production of carotenoids were up-regulated compared to control plants transferred for the same time to NL (Fig. 3). *PSY3* transcripts were undetectable in leaves from NL or HL samples (Fig. 3A), whereas chlorophylls remained virtually unchanged (Fig. 3B). The increase in carotenoid levels associated to HL exposure of WT plants was significantly repressed in *psy2* mutants and attenuated in *psy1* mutants (Fig. 3B). The potential photosynthetic capacity estimated from the measurement of the maximum quantum yield of photosystem II (Fv/Fm) was reduced in leaves from the two *psy2* alleles under normal conditions (Fig. 3C), similar to that observed for ΦPSII (Fig. 2C). Upon transfer from NL to HL, Fv/Fm progressively decreased in both WT and PSY-defective mutants, but the drop was stronger in *psy1* mutants and highest in *psy2* alleles (Fig. 3C). NPQ was also reduced in HL-exposed *psy1* and *psy2* mutants compared to WT controls, with *psy2* plants showing lower values than *psy1* alleles (Fig. 3D). These results suggest a main role for PSY2 and a supporting role for PSY1 in supplying phytoene when enhanced carotenoid synthesis is needed for photoprotection.

**Fig. 3.**
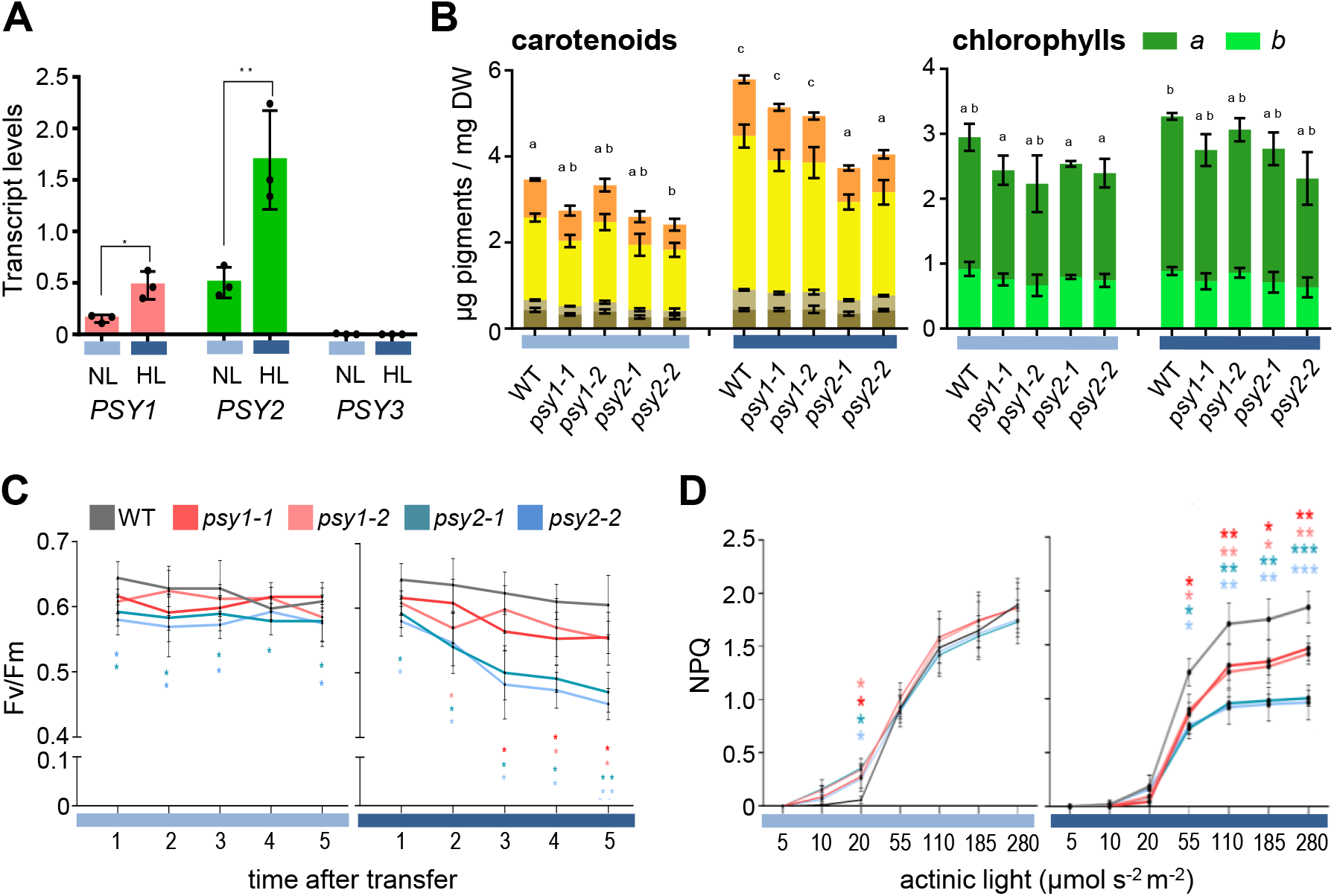
PSY2 is the main isoform required for photoprotection. Tomato WT and mutant seedlings germinated and grown under normal light conditions were left for 5 more days under the same light conditions (NL; pale blue) or transferred to high light (HL; dark blue) for the same time. **(A)** RT-qPCR analysis of *PSY1, PSY2* and *PSY3* transcript levels in WT seedlings at the end of the experiment normalized using the *ACT4* gene. Data correspond to mean and SD of n=3 independent biological replicates. Asterisks indicate statistically significant differences between means relative to NL conditions (*t*-test). **(B)** Total carotenoid and chlorophyll levels in WT and mutant seedlings exposed to either NL or HL. In the carotenoid plot, colors correspond to the species shown in Fig. 1A. Mean and SD of n=3 independent biological replicates are shown. Bar letters represent statistically significant differences (*P* < 0.05) among means according to post hoc Tukey’s tests run when one way ANOVA detected different means. **(C)** Maximum quantum yield of photosystem II (Fv/Fm) during the indicated treatments. **(D)** Non-photochemical quenching (NPQ) values at the indicated times of exposure to either NL or HL upon increasing actinic light. In (C) and (D), values represent the mean and SD of four different leaf areas from three different seedlings and asterisks indicate statistically significant differences among means in each differential time point (one way ANOVA followed by Tukey’s test). *, *P* < 0.05; **, P < 0.01, *** *P* < 0.001).

### PSY1 is supported by PSY2 to produce carotenoids for flower and fruit pigmentation

Besides their essential role in chloroplasts, carotenoids accumulate in specialized plastids named chromoplasts that provide distinctive yellow, orange and red colors to non-photosynthetic tissues such as flower petals and ripe fruit. In tomato, carotenoids (mainly conjugated xanthophylls) are responsible for the yellow color of flower petals (Fig. 4) (Ariizumi et al., 2014). As previously reported for PSY1-defective lines (Bird et al., 1991; Bramley et al., 1992; Fraser et al., 1999), *psy1-1* and *psy1-2* alleles showed flowers of a paler yellow color than the WT (Fig. 4A and Fig. S4). HPLC analysis of free and conjugated xanthophyll content showed a reduction of about 50% in PSY1-defective compared to WT corollas (Fig. 4B). While the absence of *PSY3* transcripts in flowers (Fig. S6) suggests that PSY2 feeds the production of the carotenoids detected in PSY1-defective fruit, a reduction of PSY2 activity in *psy1-2* mutants with the *psy2-1* mutation in heterozygosis, herein referred to as *psy1 PSY2(+/-)*, resulted in only a marginal reduction in carotenoid levels compared to *psy1-2* flowers (Fig. 4B). Both *psy2-1* and *psy2-2* mutant alleles showed normal-looking flowers (Fig. 4A) with a virtually WT carotenoid profile (Fig. 4B), but carotenoid levels were slightly reduced in flowers of *psy2 PSY1(+/-)* plants with only one *PSY1* gene copy in a *psy2-1* background (Fig. 4B). We therefore conclude that an excess of PSY1 activity ensures enough carotenoid production in tomato flower corollas.

**Fig. 4.**
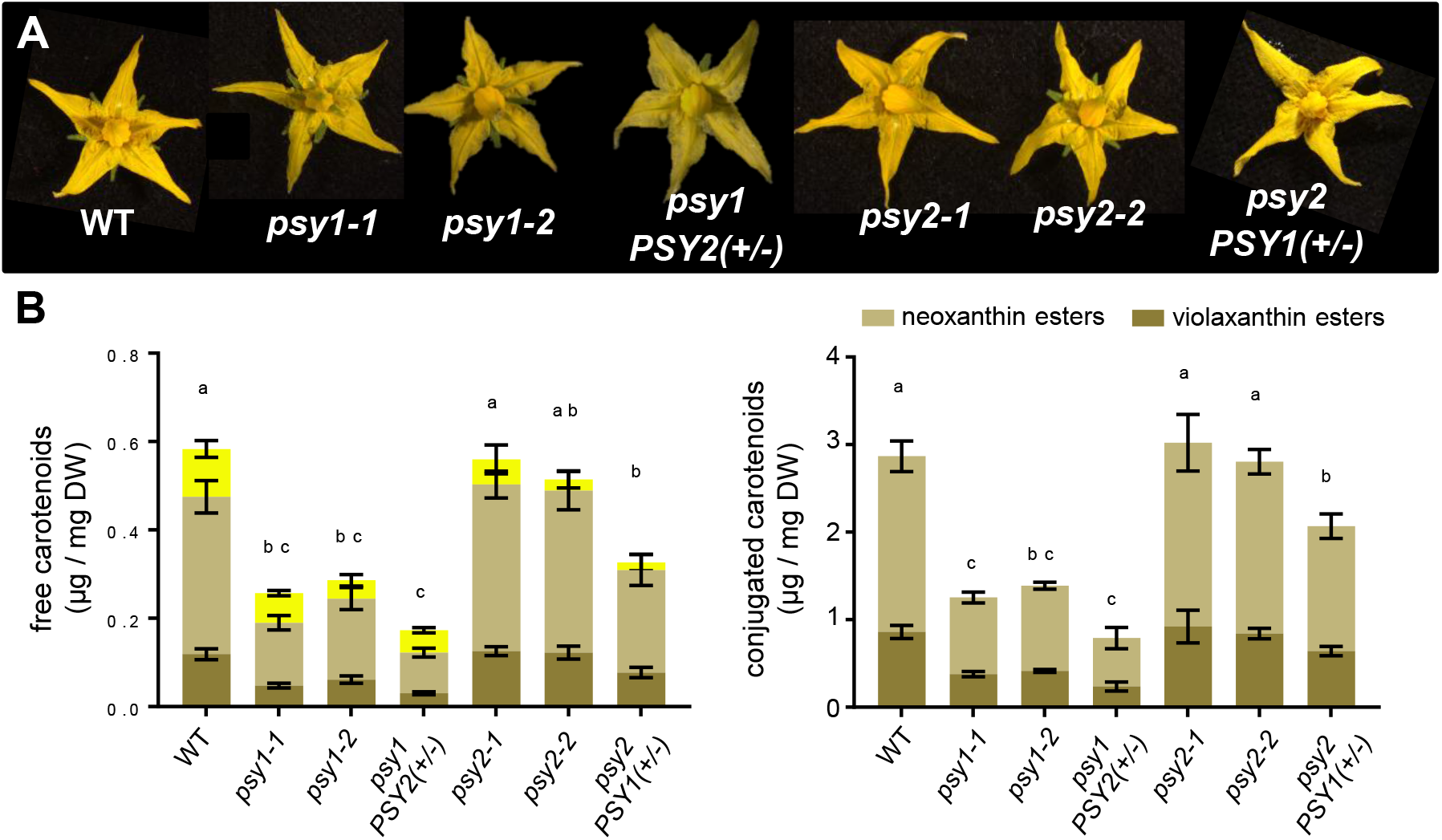
PSY1 is the main isoform contributing to carotenoid biosynthesis in petal chromoplasts. **(A)** Representative images of anthesis (fully open) flowers of the indicated lines. **(B)** Total levels of free and conjugated carotenoids in petals. In the free carotenoid plot, colors correspond to the species shown in Fig. 1A. Mean and SD of n=3 independent biological replicates are shown. Bar letters represent statistically significant differences (*P* < 0.05) among means according to one way ANOVA followed by post hoc Tukey’s tests.

The most characteristic phenotype of PSY1-defective tomato lines is the yellow color of the ripe fruit (Bird et al. 1991; D’Ambrosio et al., 2018; Fraser et al., 1999; Fray & Grierson, 1993; Gupta et al., 2022; Kachanovsky et al., 2012; Kang et al., 2014; Karniel et al., 2022). Tomato fruit ripening is a carotenoid-demanding process as great amounts of lycopene and, to a lower extent, β-carotene are produced to provide the characteristic red and orange color to the ripe fruit flesh: the pericarp (Fig. 5). Besides carotenoid synthesis, ripening also involves degradation of chlorophylls after the fruit reaches its final size at the mature green (MG) stage, which changes the fruit color from the breaker (B) stage (Fig. 5A). Previous reports have shown that loss of PSY1 activity does not impact carotenoid levels at the MG stage but it results in a drastic reduction in pericarp carotenoid levels in ripe fruit, which show a yellowish color due to flavonoid compounds such as naringenin chalcone (D’Ambrosio et al., 2018; Fraser et al., 1999; Fray & Grierson, 1993; Kachanovsky et al., 2012; Kang et al., 2014). Consistently, our edited lines with reduced PSY1 levels showed WT carotenoid and chlorophyll levels in the pericarp of MG fruit (Fig. 5B). Also as expected, analysis of pericarp carotenoid contents at six days after the B stage (B+6) showed extremely low (but still detectable) levels of carotenoids (lutein and β-carotene) in PSY1-defective fruit (Fig. 5C). To investigate the contribution of PSY2 to the residual carotenoid contents of B+6 (i.e. ripe) fruit with a complete loss of PSY1, we compared the carotenoid profile of *psy1-2* and *psy1 PSY2(+/-)* fruit. A reduction in total carotenoids was observed in *psy1-2 PSY2(+/-)* relative to *psy1-2* fruit (Fig. 5C) but it was only statistically significant for β-carotene. In agreement with the conclusion that PSY1 is by far the main contributor to carotenoid production in the pericarp of ripe fruit, complete loss of PSY2 in single *psy2-1* mutant fruit had no impact in carotenoid levels compared to WT fruit whereas a statistically significant reduction of pigment contents was found when PSY1 activity was genetically reduced in *psy2 PSY1(+/-)* fruit (Fig. 5C).

**Fig. 5.**
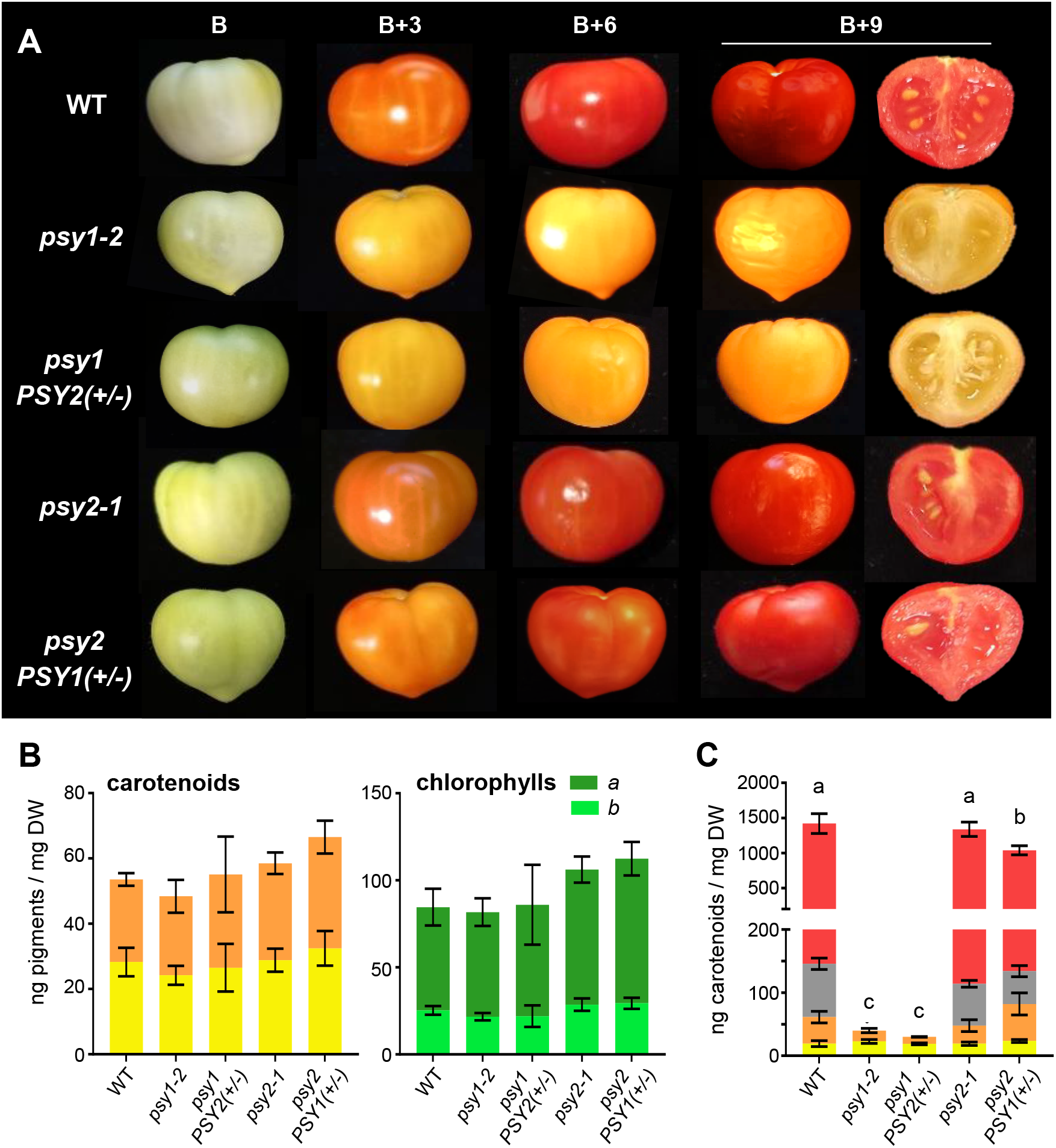
PSY1 is the main isoform contributing to carotenoid biosynthesis in fruit pericarp chromoplasts. **(A)** Representative images of WT and mutant fruit collected at the breaker (B) stage and left to ripe off-vine in a controlled environment chamber for the indicated times (in days). **(B)** Total carotenoid and chlorophyll levels in WT and mutant in the pericarp of fruit collected from the plants at the MG stage. **(C)** Total carotenoid levels in WT and mutant in the pericarp of fruit collected from the plants at the B+6 stage. In the carotenoid plots, colors correspond to the species shown in Fig. 1A. In all the plots, mean and SD of n=3 independent biological replicates are shown. Bar letters represent statistically significant differences (*P* < 0.05) among means according to post hoc Tukey’s tests run when one way ANOVA detected different means.

After the B stage, our *psy1*-*2* and *psy1 PSY2(+/-)* fruits acquired a distinctive yellowish color but PSY2-defective *psy2-1* and *psy2 PSY1(+/-)* fruits were undistinguishable from WT fruits (Fig. 5A). Color analysis using TomatoAnalyzer showed that color changes in *psy2-1* and *psy2-2* fruits occurred at a similar rate as in WT controls (Fig. 6A). To test whether mutant fruit showed other ripening-associated phenotypes besides color, the expression of ripening marker genes such as *E8* (Solyc09g089580) and *ACS2* (Solyc01g095080) was quantified by RT-qPCR (Barja et al., 2021). As shown in Fig. 6B, the expression profile of these genes was very similar in WT and *psy2-1* fruit during ripening. By contrast, the peak of *E8* and *ACS2* expression observed at the B stage was significantly reduced in *psy1-2* fruit (Fig. 6B).

**Fig. 6.**
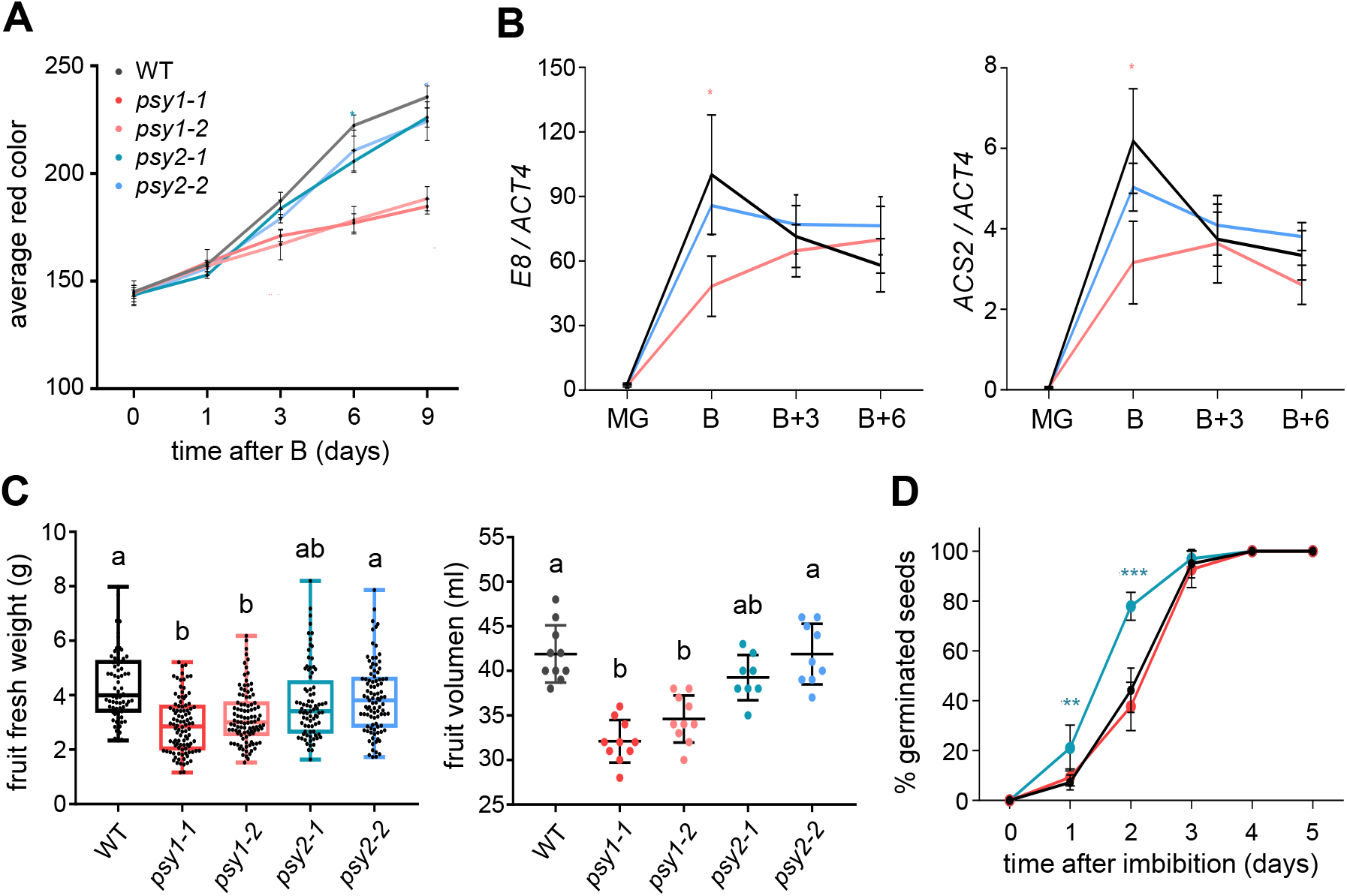
Fruit development and seed germination are differentially impacted in mutants defective in PSY1 or PSY2. **(A)** Average red color quantification (arbitrary units) of fruit collected at the B stage and left to ripe off-vine in chambers for the indicated times. Values represent the mean and SD of n=3 different fruits for each time point. **(B)** RT-qPCR analysis of *E8* and *ACS2* transcript levels in WT and mutant fruit collected from the plant at the indicated stage. Data correspond to mean and SD of n=3 independent biological replicates. Asterisks indicate statistically significant differences relative to the WT (*t*-test, *P* < 0.05). **(C)** Weight and volume of fully ripe fruits of the indicated genotypes. In the boxplot, the lower and upper boundary of the boxes indicate the 25th and 75th percentile, respectively; the line inside the boxes represent the median; dots mark individual data values; and whiskers above and below the boxes indicate the maximum and minimum values. In the dot plots, central line represents the mean and whiskers represent SD. Different letters represent statistically significant differences (one way ANOVA followed by Tukey’s multiple comparisons test, *P* < 0.05). **(D)** Kinetics of germination of WT and mutant seeds after imbibition. Error bars indicate SD of n=6 biological replicates with 25 seeds each. Asterisks indicate statistically significant differences among means relative to WT samples (*t*-test: **, *P* < 0.01, *** *P* < 0.001).

### PSY1 and PSY2 are major contributors to ABA synthesis in tomato fruit pericarp and seeds, respectively

ABA is a carotenoid-derived phytohormone (Fig. 1A) which, besides regulating plant adaptation to abiotic stress conditions and promoting seed dormancy, appears to regulate fruit growth and development in tomato (Leng et al., 2014; Nambara & Marion-Poll, 2005; Zhang et al., 2009). Indeed, reduced hormone levels in mutants defective in ABA biosynthetic genes such as *notabilis* (*NOT/NCED*), *sitiens* (*SIT/AAO3*), and *flacca* (*FLC/ABA3*) are associated to slower ripening but also to reduced fruit size and accelerated seed germination (De Castro & Hilhorst 2006; Galpaz et al., 2008; Groot & Karssen, 1992; McQuinn et al., 2020; Nitsch et al., 2012). ABA levels in pericarp and seeds peak around the B stage, preceding the burst of ethylene biosynthesis that regulates many aspects of the ripening process in a climacteric fruit such as tomato (Berry & Bewley 1992; De Castro & Hilhorst 2006; Diretto et al., 2020; Zhang et al., 2009). Quantification of ABA in pericarp and seed samples from WT and mutant fruit at the B stage showed decreased levels of the hormone in the pericarp of *psy1-2* fruit and the seeds of *psy2-1* samples (Fig. 7). Consistent with the decrease in pericarp ABA levels, fruits lacking PSY1 not only showed a reduced peak of ripening-related gene expression (Fig. 6B) but also lower fruit weight and volume compared to WT and PSY2-defective fruits (Fig. 6C). We also analyzed the germination (root emergence) of WT and mutant seeds freshly collected from ripe fruits. Accordingly to the reduced levels of ABA in the seeds of PSY2-defective mutants (Fig. 7), *psy2-1* seeds showed an accelerated germination compared to WT and *psy1-2* seeds (Fig. 6D).

**Fig. 7.**
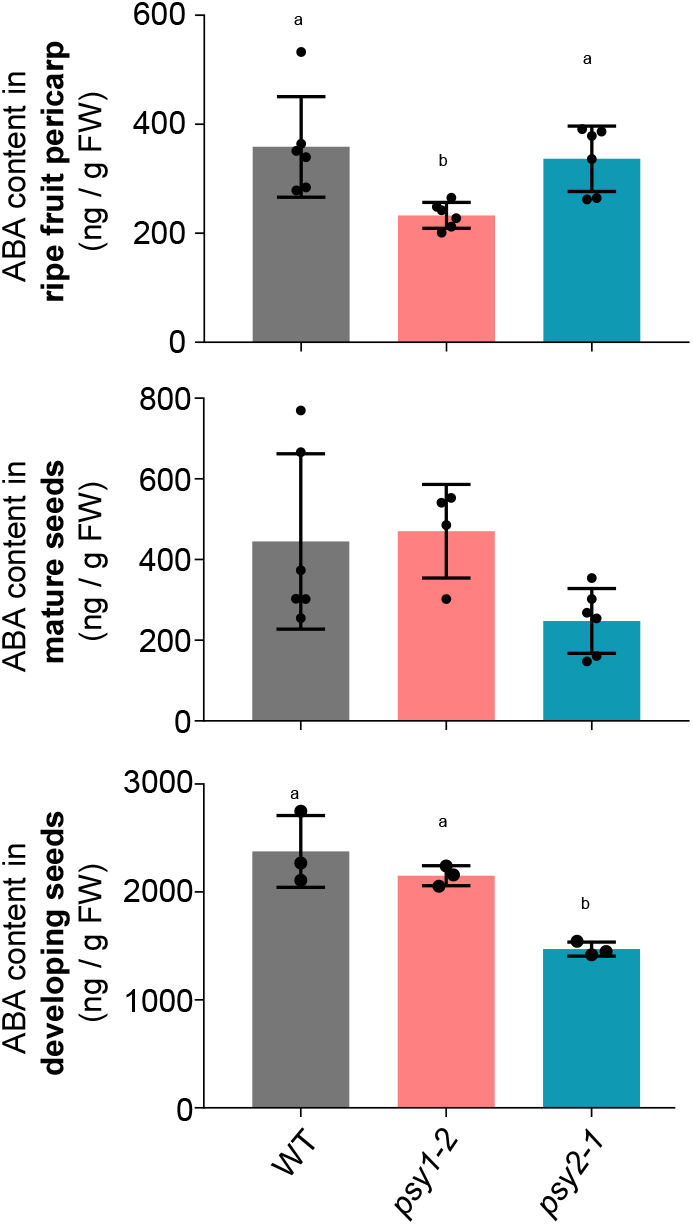
ABA levels are different in fruit pericarp and seed samples from mutants defective in PSY1 or PSY2. Pericarp and mature seed samples were collected from ripe (B+6) fruit, whereas developing seeds were collected from immature fruits. Values correspond to the mean and SD of samples collected from n≥3 independent fruits. Different letters represent statistically significant differences among means (one way ANOVA followed by Tukey’s multiple comparisons test, *P* < 0.05).

The described data suggest that PSY1 might be most important for ABA production in the pericarp and PSY2 in seeds. This conclusion is only partially consistent with transcript abundance profiles during fruit pericarp and seed development (Fig. S6-S8). In the pericarp, *PSY1* is expressed at higher levels than *PSY2* from early stages of fruit development, and the differences become much more dramatic after the MG stage (Fig. S6 and S7). In developing seeds, both genes are expressed at similar levels (Fig. S6 and S8) and yet ABA contents are reduced in the *psy2-1* mutant but not in the *psy1-1* line when compared to the WT (Fig. 7). As fruits ripe, *PSY1* expression increases and *PSY2* expression decreases in mature seeds (Fig. S6 and S8), which similar to developing seeds only show reduced ABA levels when PSY2 activity is removed (Fig. 7). To provide further evidence on the role of specific PSY isoforms in the production of ABA involved in the control of seed dormancy, we blocked carotenoid production in all the tissues of MG fruits of WT and mutant fruits using specific inhibitors (Fig. 8). Specifically, we used the MEP pathway inhibitor fosmidomycin (FSM) and the phytoene desaturase inhibitor norflurazon (NFZ) (Fig. 1A). WT, *psy1-2* and *psy2-1* fruits were collected from the plant at the MG stage and injected with one of the inhibitors or a mock solution (water). After twelve days, WT and *psy2-1* fruits treated with either FSM or NFZ showed a yellow color identical to that of *psy1-2* fruit treated with mock or inhibitor solutions (Fig. 8A), confirming that both FSM and NFZ successfully inhibited carotenoid production, at least in the pericarp. At this point, seeds were collected from the detached fruits, dried overnight, and immediately used for germination assays (Fig. 8B). In the case of WT and *psy1-2* seeds, germination was accelerated by the treatment with either FSM or NFZ, suggesting an inhibitor-mediated blockage of carotenoid and hence downstream ABA production in seeds. By contrast, inhibitor treatment had no effect on the germination rate of *psy2-1* seeds (Fig. 8B). These results support the conclusion that seed dormancy is independent of the ABA content of the fruit pericarp or developing seeds but it is regulated by ABA produced in mature seeds from PSY2-derived carotenoids.

**Fig. 8.**
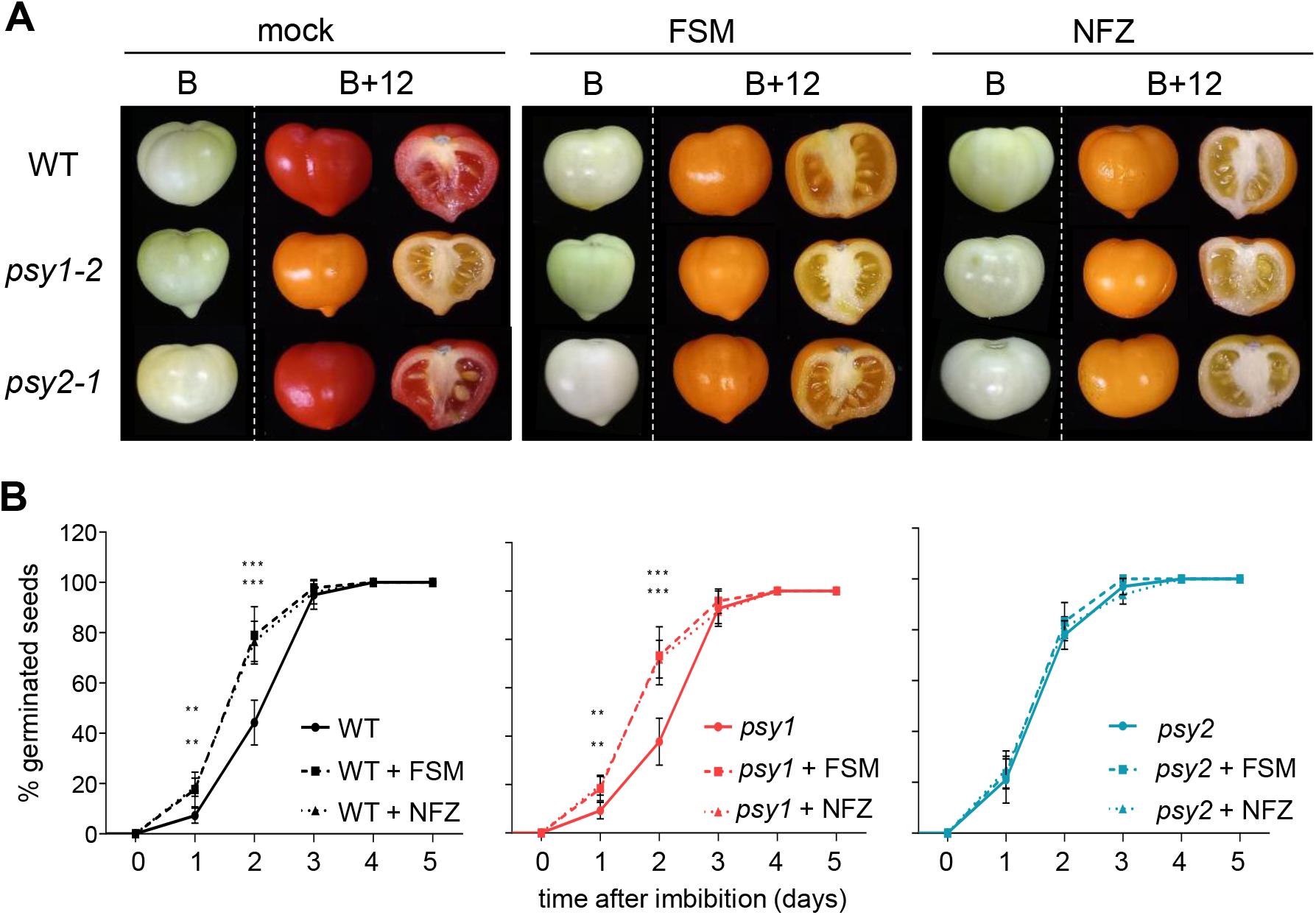
Germination of tomato seeds is regulated by ABA produced in mature seeds from PSY2-derived carotenoids. **(A)** Representative images of WT and mutant fruits treated with fosmidomycin (FSM), norflurazon (NFZ) or a mock solution at the MG stage and then allowed to ripe off-vine. **(B)** Kinetics of germination of fresh seeds collected from fruits as those shown in (A) at the B+12 stage. Error bars indicate SD of n=6 biological replicates with 25 seeds each. Asterisks indicate statistically significant differences among means relative to WT samples (*t*-test: **, *P* < 0.01, *** *P* < 0.001).

## Discussion

PSY catalyzes the first committed and main rate-determining step of the carotenoid pathway. In most plants, several PSY isoforms control the production for carotenoids in different tissues and in response to developmental or environmental cues that require an enhanced production of these photoprotective pigments (Zhou et al., 2022). The presence of three PSY isoforms in tomato has been known for a long time, but genetic evidence on their physiological roles was only available for PSY1. Removal of PSY1 activity in mutants or silenced lines leads to strongly reduced levels of carotenoid pigments in ripe fruit and, to a lower extent, in corollas but unchanged carotenoid levels in green tissues, which led to conclude that PSY1 is mainly involved in carotenoid biosynthesis in chromoplasts (Bird et al. 1991; Bramley et al., 1992.; D’Ambrosio et al., 2018; Fantini et al., 2013; Fraser et al., 1999; Giorio et al., 2008; Kang et al., 2014). Compared to *PSY1, PSY2* expression is higher in leaves and increases more strongly during seedling deetiolation, supporting the conclusion that PSY2 might be the main isoform producing phytoene for carotenoids involved in photosynthesis and photoprotection (Barja et al., 2021; Bartley & Scolnik, 1993; Fraser et al., 1999; Gupta et al., 2022) (Fig. S6). *PSY3* expression levels are very low in all the tissues compared to *PSY1* and *PSY2* (Giorio et al., 2008; Stauder et al., 2018) (Fig. S6). Similar to most members of the PSY3 clade, tomato *PSY3* expression is highest in roots, where it is induced during arbuscular mycorrhizal (AM) fungi colonization (Barja et al., 2021; Stauder et al., 2018; Walter et al., 2015). Based on these expression data, it was concluded that PSY3 might have a main role in roots, supplying phytoene to produce carotenoids and derived SLs and apocarotenoid molecules essential for the establishment of the AM symbiosis (Baslam et al., 2013; Fester et al., 2002; Ruiz-Lozano et al., 2016; Stauder et al., 2018). This work aimed to genetically test the hypothesis that besides the main role of PSY1 for carotenoid production in flowers and fruit (chromoplasts), PSY2 in green tissues (chloroplasts) and PSY3 in roots (leucoplasts), tomato PSY isoforms might also provide extra phytoene when a sudden requirement of carotenoid production could not be met by the isoform normally operating in a particular tissue. The generation of lines defective in PSY1 and/or PSY2 reported here provided strong genetic support to correctly frame this conclusion and it went a step beyond by unveiling a role for particular PSY isoforms in tissue-specific ABA production.

Complete loss of PSY activity in Arabidopsis results in albino seedlings (Pokhilko et al., 2015). In *Nicotiana benthamiana* (a closer relative to tomato), several genes encode PSY1, PSY2 and PSY3 homologues, but the virus-induced silencing of only those for PSY1 and PSY2 results in leaf bleaching, lower carotenoid levels and reduced photosynthetic parameters such as ΦPSII, Fv/Fm and NPQ (Wang et al., 2021). Similarly, we observed a seedling-lethal albino phenotype in tomato lines lacking PSY1 and PSY2 but retaining a functional *PSY3* gene (Fig. 1C). This result demonstrates that PSY3 is unable to produce enough phytoene to support photosynthetic shoot development when PSY1 and PSY2 activities are missing. Indirectly, the result also provides genetic evidence supporting a root-restricted role for tomato PSY3. In the shoot, both PSY1 and PSY2 appear to provide precursors for carotenoid biosynthesis in chloroplasts under normal growth conditions (Fig. 2). However, the lower up-regulation of *PSY1* expression compared to *PSY2* in response to HL (Fig. 3A) together with the reduced impact of the loss of PSY1 function on carotenoid levels and photosynthetic performance (Fig. 2 and 3) supports the model of a predominant role for PSY2 and a supporting contribution of PSY1 to carotenoid biosynthesis in tomato chloroplasts for photoprotection.

Besides chloroplasts, tomato plants accumulate very high levels of carotenoids in the chromoplasts that develop in flower corollas and ripening fruit pericarp. Loss of PSY1 had a much stronger impact than removing PSY2 on total carotenoid levels of both tissues. However, the effect in flowers (Fig. 4) was much less dramatic than in fruit (Fig. 5). Despite *PSY2* is highly expressed in petals (Giorio et al., 2007) (Fig. S6) and PSY2 catalytic activity appears to be higher than that of PSY1 (Cao et al., 2019), complete absence of PSY2 had no effect of petal carotenoids (Fig. 4). By contrast, a 50% decrease compared to the WT was observed in PSY1-defective corollas (Fig. 4), suggesting that PSY2 only produces phytoene for carotenoid synthesis in flower chromoplasts when PSY1 activity is missing. A similar conclusion was deduced for PSY2 during fruit ripening (Gupta et al., 2022; Karniel et al., 2022). In fruit pericarp tissues, carotenoid levels were unaffected in mutants defective in PSY1 and PSY2 until the onset of ripening (Fig. 5), supporting our conclusion that loss of one of the two isoforms can be rescued by the remaining isoform in chloroplasts. As chloroplasts differentiate into chromoplasts, however, the contribution of PSY1 to the production of pericarp carotenoids becomes much more predominant, mainly supported by a dramatic up-regulation of gene expression (Cao et al., 2019; Giorio et al., 2007) (Fig. S4 and S6). Without PSY1, ripe fruit accumulate a very small amount of carotenoid pigments (Fig. 5C). These carotenoids (mainly lutein and β-carotene) might be remnants of the carotenoids present in MG fruits. However, PSY activity has been measured in chromoplasts of PSY1-defective fruit (Fraser et al., 1999), and a PSY2-dependent increase in carotenoid synthesis was observed during ripening of PSY1-lacking fruit treated with different inhibitors (Gupta et al., 2022; Karniel et al., 2022). A role for PSY2 in the production of β-carotene in pericarp chromoplasts during ripening can also be deduced from the reduced accumulation of this carotenoid in *psy1 PSY2(+/-)* compared to *psy1-2* fruit (Fig. 5C). A similar role distribution has been recently described in pepper (*Capsicum annuum*), where PSY1 is the isoform supporting the bulk of pericarp carotenoid biosynthesis during fruit ripening and PSY2 is mainly associated to chloroplast-containing tissues (leaves, stems) but it also contributes to produce carotenoids in fruit chromoplasts (Jang et al., 2020; Wei et al., 2021). It has been proposed that recruitment of primary (i.e. photosynthetic) carotenoids as secondary metabolites for flower and fruit pigmentation likely required duplication and further subfunctionalization of genes encoding rate-controlling steps, including PSY (Galpaz et al., 2006; Giorio et al., 2008). In tomato, the duplicated pathway might have been originally employed for flower pigmentation and later for fruit pigmentation, explaining why all tomato species have yellow flowers but only some develop fruit chromoplasts (Galpaz et al., 2006; Giorio et al., 2008).

Besides providing strong genetic evidence supporting long-standing models on the subfunctionalization of tomato PSY1 and PSY2 isoforms to feed the carotenoid pathway in particular tissues, our results have unveiled isoform-specific roles in ABA-regulated processes in tomato fruit and seeds. Genetic and pharmacological interference with carotenoid biosynthesis was previously shown to impact ABA-regulated characters such as fruit size, the onset of fruit ripening and seed dormancy (Diretto et al., 2020; Galpaz et al., 2008; McQuinn et al., 2020; Zhang et al., 2009). Also, pharmacological approaches had provided evidence suggesting (but not demonstrating) that PSY2 might be involved in the production of ABA in tomato fruits (Gupta et al., 2022), most particularly in seeds (Rodriguez-Concepcion et al. 2001). Here we showed that in the absence of PSY1, PSY2-derived carotenoids sustain the production of about 2/3 of the ABA measured in the pericarp of tomato B fruit (Fig. 7). The 1/3 reduction was sufficient to trigger phenotypes associated to low ABA levels in *psy1* fruit, including an attenuated expression of ethylene-associated ripening gene (Fig. 6B) and a lower fruit weight and volume (Fig. 6C), suggesting that a threshold of ABA is required to support normal fruit growth and ripening. Alternatively, PSY1-derived carotenoids might be responsible for the production of ABA in specific tissues or cell compartments causing the observed phenotypes. A differential channeling of phytoene produced by either PSY1 or PSY2 to produce carotenoids for specific ABA pools is supported by the seed germination experiments. When PSY2 is not present, PSY1 still produces about 2/3 of the ABA measured in developing and mature seeds (Fig. 7) but this relatively high amount of remaining ABA is not enough to prevent a germination delay phenotype in the *psy2* mutant (Fig. 6D). Furthermore, complete block of carotenoid (and hence downstream) ABA production in MG fruit with inhibitors did not exacerbate the seed dormancy phenotype of the *psy2* mutant (Fig. 8B). These results strongly suggest that only PSY2-derived carotenoids produced after the MG stage are used to generate the ABA that regulates dormancy in tomato seeds. Following an initial phase of tissue differentiation, tomato seed development proceeds as fruit expand with a second phase that includes the accumulation of nutrient reserves and the acquisition of germination and desiccation tolerance (De Castro & Hilhorst 2006) (Fig. S8). When fruits reach their final size at the MG stage, seeds achieve show full germinability. Later, as fruit start to ripe, ABA production peaks and mature seeds dry and acquire their dormancy (De Castro & Hilhorst 2006). This transient accumulation of ABA is mainly supplied by the embryo (Berry & Bewley 1992). Our results confirm that ABA produced by PSY2 in seeds during ripening regulates seed dormancy (Fig. 8). Strikingly, the contribution of PSY2 to produce this ABA could not be predicted based on available expression data. Thus, *PSY2* expression is higher in developing seeds but then it drops from the MG stage in embryonic and other tissues of mature seeds (Fig. S8). While a *PSY2*-like profile is observed for the *NOT/NCED* gene (Solyc07g056570), which encodes the first enzyme specific for ABA biosynthesis (Fig. S8), downstream genes of the pathway such as *SIT/AAO3* (Solyc01g009230) and *FLC/ABA3* (Solyc07g066480) are more highly expressed in mature seeds (including embryos) (Fig. S8). Although *PSY1* expression is also higher in mature seeds, little to no expression was found in embryos. These results clearly illustrate the challenges of deducing function based only on gene expression profiles.

A question arising from our data is how interference with PSY activity is specifically translated into changes in the production of ABA (Fig. 1A and Fig. S8). A possible scenario would be the existence of metabolons channeling GGPP to ABA in cells from the pericarp or the seed. GGPP required to produce pericarp carotenoids and ABA during fruit ripening is mainly supplied by the GGPPS isoform SlG3 with a supporting contribution of SlG2 (Barja et al., 2021). While SlG2 can interact with both PSY1 and PSY2, no interaction was reported for SlG3 in transient co-expression assays in *N. benthamiana* leaves (Barja et al., 2021). It is possible that interaction of SlG3 with particular PSY isoforms requires specific partners only found in tomato pericarp (to interact with PSY1) or seed (to interact with PSY2) tissues. In agreement with the existence of metabolons or any other kind of metabolic channeling, the extremely low PSY2 activity present in the pericarp of *psy1* fruit appears to be more directly involved in the production of the β-carotene instead of lutein (Fig. 5C), i.e. it might be preferentially acting to produce carotenoids that could then be used as precursors for ABA synthesis (Fig. 1A). However, the channeling of specific pools of carotenoids all the way to ABA is harder to fit in a metabolon-dependent model due to the large number of reactions and the diversity of subcellular localizations reported for the enzymes involved, which include several cytosolic steps following the cleavage of β-carotene-derived xanthophyll precursors (Nambara & Marion-Poll, 2005) (Fig. 1A and Fig. S8). Alternatively, expression of specific isoforms in particular tissue microdomains controlling fruit ripening (directly or indirectly through ethylene), pericarp growth, or seed dormancy might explain why only PSY1-derived ABA appears to contribute to fruit ripening and only PSY2-derived ABA influences seed germination.

In summary, we show that both PSY1 and PSY2 support carotenoid production in tomato shoots with diverging contributions in different tissues: PSY2 > PSY1 in leaves (i.e. chloroplasts), PSY1 > PSY2 in corollas, and PSY1 >> PSY2 in fruit pericarp tissues. Furthermore, we demonstrate a differential contribution to the production of ABA of PSY1 in the pericarp (to regulate fruit growth and ripening) and PSY2 in the seeds (to control dormancy). Further work should determine the mechanism by which the production of phytoene by given PSY isoforms is eventually channeled to produce ABA is particular locations for specific functions.

## Materials and methods

### Plant material, treatments and sample collection

Tomato (*Solanum lycopersicum* var. MicroTom) plants were used for all the experiments. Seeds were surface-sterilized by a 30 min water wash followed by a 15 min incubation in 10 ml of 40% bleach with 10 µl of Tween-20. After three consecutive 10 min washes with sterile milli-Q water, seeds were germinated on plates with solid 0.5x Murashige and Skoog (MS) medium containing 1% agar (without vitamins or sucrose). The medium was supplemented with kanamycin (100 μg/ml) when required to select transgenic plants. Plates were incubated in a climate-controlled growth chamber (Ibercex) at 26°C with a photoperiod of 14 h of white light (photon flux density of 50 μmol m^−2^ s^−1^) and 10 h of darkness. After 10-14 days, seedlings were transferred to soil and grown under standard greenhouse conditions (14 h light at 25 ± 1 °C and 10 h dark at 22 ± 1 °C). Young leaves were collected from 4-week-old plants and they correspond to growing leaflets from the fourth and fifth true leaves. Petal samples were collected from anthesis flowers. Fruit pericarp samples were collected at different stages, including mature green (MG, about 30 days post-anthesis), breaker (B, 2-3 days later, when the first symptoms of chlorophyll degradation and carotenoid accumulation became visually obvious), and several days after breaker. After collection, samples were immediately frozen in liquid nitrogen and stored at -80°C. For fruit weight determination, 100 fully ripe individual fruits from each genotype were collected and weighted one by one using a precision scale (Kern). Fruit volume was estimated in 10 pools of 10 fruits each by measuring the displaced water volume in a graduated cylinder. For inhibitor treatments, MG fruits were collected from the plant and measured to estimate their volume. Then, a Hamilton syringe was used to inject 2-5 µl of sterile water or inhibitor solution into the fruit. The exact volume of fosmidomycin (FSM, Sigma) or norflurazon (NFZ, Zorial, Syngenta) solution to inject was calculated based on the fruit volume so the final concentration in the fruits was 200 µM FSM or 50 µM NFZ. After injection, fruits were kept in a climate-controlled growth chamber at 26°C for 12 days and then seeds were collected and immediately used for germination assays on 0.5x MS plates. Germination was scored based on root protrusion.

### Generation of CRISPR-Cas9 mutants and tomato transformation

For CRISPR-Cas9-mediated disruption of *PSY1* and *PSY2*, one single guide RNA (sgRNA) was designed for each gene using the online tool CRISPR-P 2.0 (Liu et al., 2017). Cloning of the CRISPR-Cas9 constructs was carried out as previously described (Barja et al., 2021) using primers listed in Table S1. As a result, a single final binary plasmid harboring the Cas9 sequence, the *NPTII* gene providing kanamycin resistance, and the sgRNAs to disrupt *PSY1* and *PSY2* was obtained and named pDE-PSY1,2 (Table S2). All constructs were confirmed by restriction mapping and DNA sequencing. *Agrobacterium tumefaciens* GV3101 strain was used to stably transform tomato MicroTom cotyledons with pDE-PSY1,2 as described (Barja et al., 2021). *In vitro* regenerated lines showing kanamycin (100 μg/ml) resistance were used for PCR amplification and sequencing of the genomic sequences. Following further segregation and PCR-based genotyping using specific primers (Table S1), stable homozygous lines lacking the Cas9-encoding transgene were obtained and named *psy1-1, psy1-2, psy2-1* and *psy2-2*. For the generation of double double mutants lacking both PSY1 and PSY2, *psy1-2* and *psy2-1* homozygous plants were crossed and the segregating F2 offspring was used for PCR-based genotyping of individual plants.

### Photosynthetic parameters

Tomato seedlings were germinated and grown for ten days under white light with a fluorescence photon flux density of 50 μmol m^−2^ s^−1^ (referred to as normal light, NL) and then either left under NL or transferred to a chamber with a more intense light of 300 μmol m^−2^ s^−1^ (referred to as high light, HL) for five more days. Chlorophyll fluorescence measurements were carried out with a Handy FluorCam (Photon Systems Instruments). Fv/Fm was measured in seedlings incubated in the dark for 30 min to allow full relaxation of photosystems. ɸPSII was measure at 30 PAR with an actinic light of 3 μmol m^−2^ s^−1^. For NPQ measurements, the following steps of actinic irradiance were used: 0, 5, 10, 20, 55, 110, 185 and 280 μmol photons m^−2^ s^−1^.

### RNA extraction and RT-qPCR analyses

Total RNA was extracted from tomato freeze-dried tissue using the PureLink RNA MINI extraction kit (Ambion). RNA was quantified using a NanoDropTM 8000 spectrophotometer (ThermoFischer Scientific) and checked for integrity by agarose gel electrophoresis. The Transcriptor First Strand cDNA Synthesis Kit (Nzytech) was used to reverse transcribe 1 μg of extracted RNA and the generated cDNA volume (20 μl) was subsequently diluted 5-fold with mili-Q water and stored at -20 °C for further analysis. Transcript abundance was evaluated via real-time quantitative PCR (RT-qPCR) in a reaction volume of 10 μl containing 2 μl of the cDNA dilution, 5 μl of SYBR Green Master Mix (Thermo Fisher Scientific), and 0.3 μM of each specific forward and reverse primer (Table S1). The RT-qPCR was carried out on a QuantStudio 3 Real-Time PCR System (Thermo Fisher Scientific) using three independent biological samples and three technical replicates of each sample. Normalized transcript abundance was calculated as previously described (Simon, 2003) using tomato *ACT4* (Solyc04g011500) as endogenous reference gene.

### Pigment quantification

Carotenoids and chlorophylls were extracted as described (Barja et al., 2021) with some modifications. Freeze-dried material from leaves (8 mg) were mixed with 375 μl of methanol as extraction solvent, 25 μl of a 10 % (w/v) solution of canthaxanthin (Sigma) in chloroform as internal control, and glass beads. Following steps were performed as described (Barja et al., 2021). Freeze-dried flower petals and fruit pericarp tissue (20 mg) were mixed in 2 ml Epperdorf tubes with 1 ml of 2:1:1 hexane:acetone:methanol as extraction solvent, 25 μl of the canthaxanthin solution, and glass beads. After vortexing the samples, 100 μl of milli-Q water were added to the mix. Then, samples were shaken for 1 min in a TissueLyser II (Qiagen) and then centrifuged at 4ºC for 5 min at maximum speed in a tabletop microfuge. The organic phase was transferred to a 1.5 ml tube and the rest was re-extracted with 1 ml of 2:1:1 hexane:acetone:methanol. The organic phases from the two rounds of extraction were mixed in the same tube and evaporated using a SpeedVac. Extracted pigments were resuspended in 200 μl of acetone by using an ultrasound bath and filtered with 0.2 μm filters into amber-colored 2 ml glass vials. Separation and quantification of individual carotenoids and chlorophylls was performed as described (Barja et al., 2021). Fruit pigmentation (Average Red Color) was measured in three different tomato fruit samples of each genotype using the default settings of the TomatoAnalyzer 4.0 software (https://vanderknaaplab.uga.edu/tomato_analyzer.html).

### Determination of ABA levels

For ABA extraction, 100 mg of frozen pericarp tissue or seeds were ground with a mortar and pestle and resuspended in a solution of 80% (v/v) methanol and 1% (v/v) acetic acid with deuterium-labelled ABA as internal standard. After shaking for 1 h at 4°C, the extract was centrifuged at maximum speed in a table top microfuge and the supertnatant was collected and dried in a SpeedVac. The dry residue was dissolved in 1% (v/v) acetic acid and run through a reverse phase column (Oasis HLB) as described (Seo et al., 2011). The eluate was dissolved in 5% (v/v) acetonitrile and 1% (v/v) acetic acid and used for UHPLC chromatography with a reverse phase 2.6 μg Accucore RP-MS column of 100 mm length x 2.1 mm i.d. (ThermoFisher Scientific). The mobile phase was 5 to 50% (v/v) acetronitrile gradient containing 0.05% (v/v) acetic acid at 400 μl/min over 21 min. Quantification of ABA was performed with a Q-Exactive mass spectrometer equipped with an Orbitrap detector (ThermoFisher Scientific) by targeted Selected Ion 100 Monitoring (SIM). The concentrations of ABA in the extracts were determined using embedded calibration curves and the TraceFinder 4.1 SP1 software.

## Acknowledgments

We thank Mª Rosa Rodriguez-Goberna for technical help with HPLC analyses and Tsuyoshi Nakagawa (Shimane University, Japan) for the Gateway vectors. We also thank members of our laboratory for helpful discussions. This work was funded by grants from Spanish MCIN/AEI/10.13039/501100011033 and European NextGeneration EU/PRTR and PRIMA programs to MR-C (PID2020-115810GB-I00 and UToPIQ-PCI2021-121941). MR-C is also supported by CSIC (202040E299) and Generalitat Valenciana (PROMETEU/2021/056). ME and EB received predoctoral fellowships from MCIN/AEI (BES-2017-080652) and Colombia’s Doctorado Exterior program (10852908329), respectively. No conflict of interest is declared.

## Author contributions

ME and MR-C designed the research; ME and EB conducted the experiments; ME, EB and MR-C analyzed and discussed data; ME and MR-C wrote the paper.

## Supplemental Figures

**Fig. S1.**
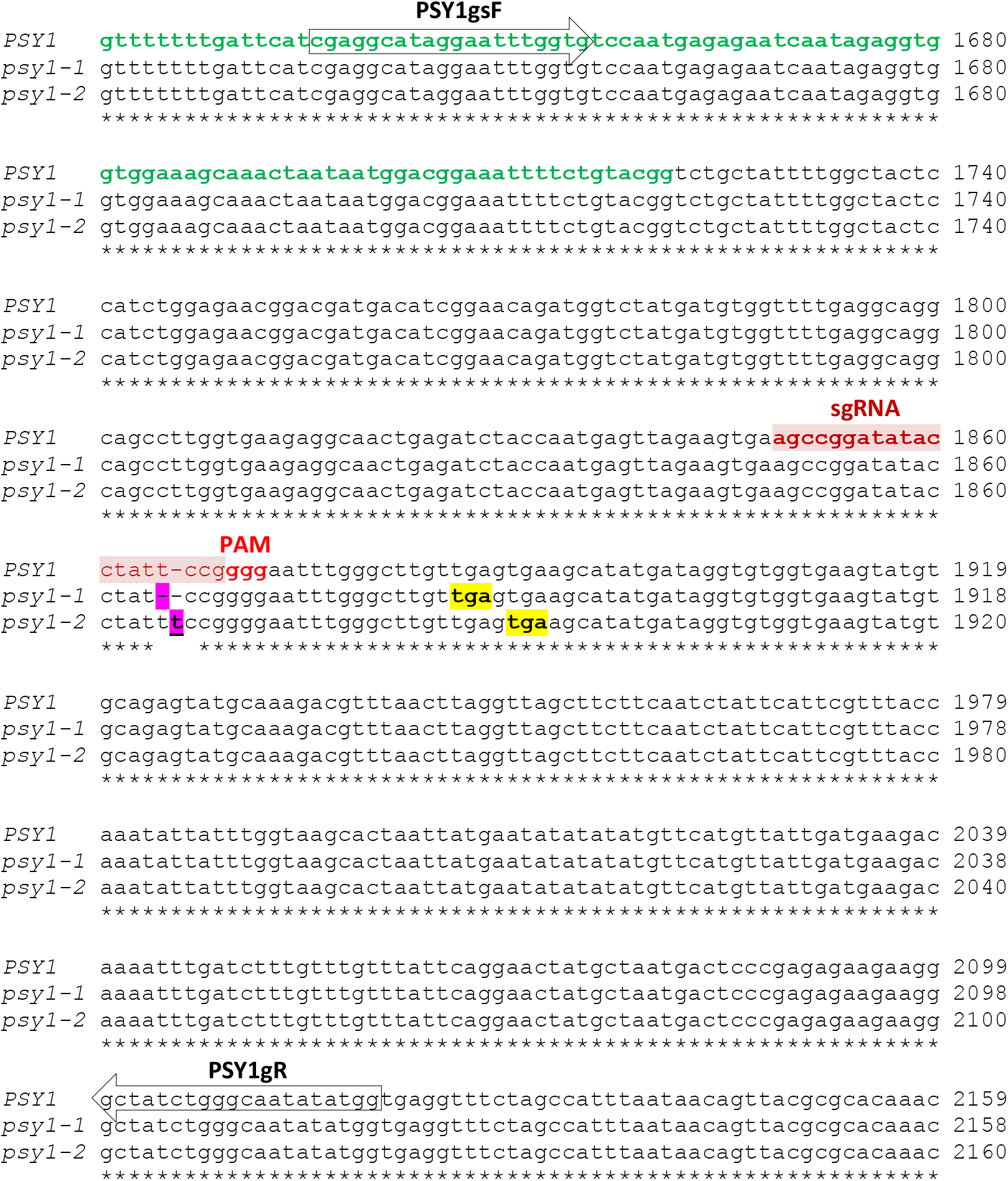
DNA sequence alignment of *PSY1* sequences from WT and CRISPR mutants. Alignment was performed using Clustal Omega with default settings (https://www.ebi.ac.uk/Tools/msa/clustalo/). The WT sequence encoding the plastid targeting sequence is marked in green and the designed single-guide RNA (sgRNA) sequence and protospacer adjacent motif (PAM) in red. The position of genotyping primers is highlighted as arrows. Mutations are boxed in pink and translation stop codons are boxed in yellow.

**Fig. S2.**
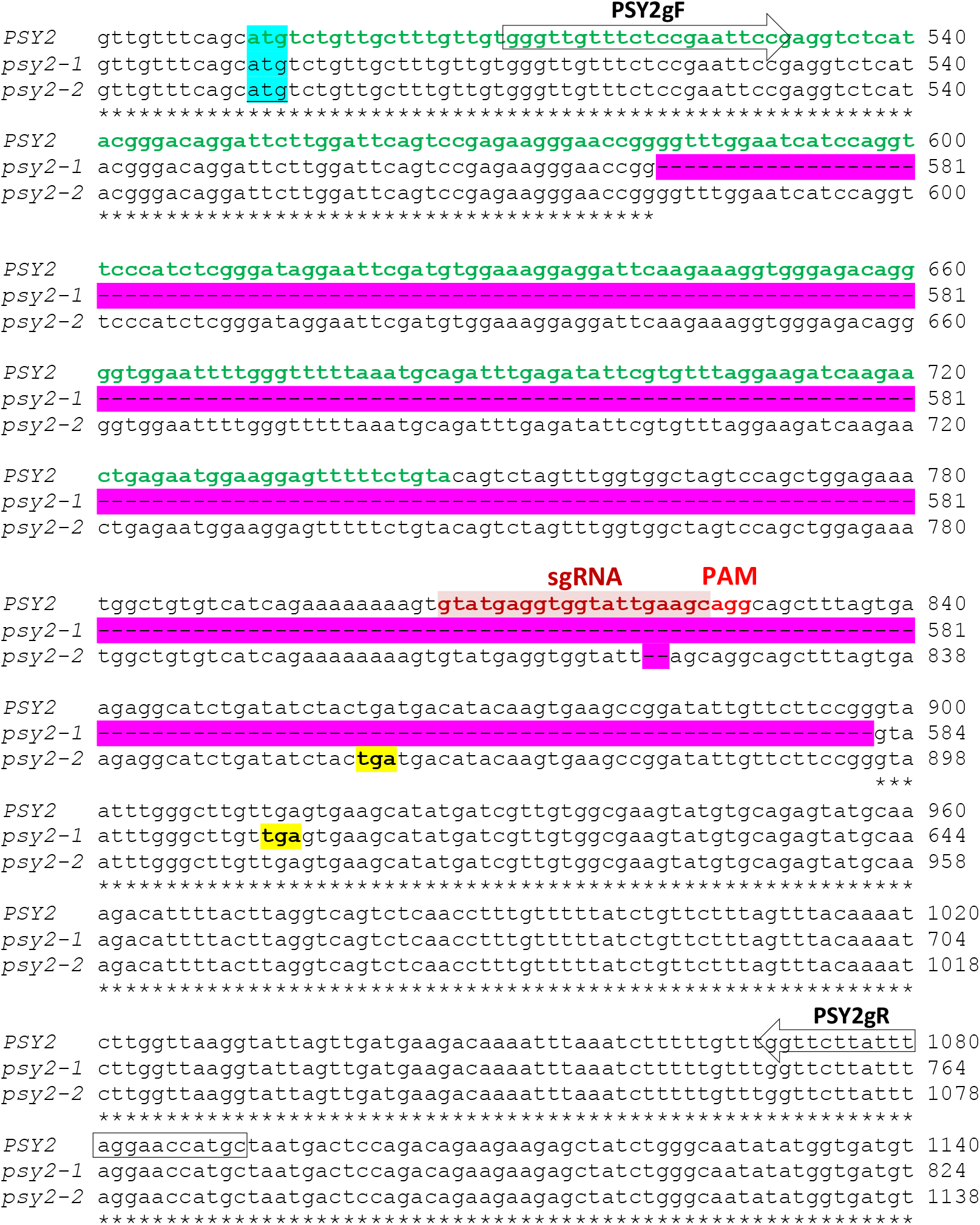
DNA sequence alignment of *PSY2* sequences from WT and CRISPR mutants. Alignment was performed using Clustal Omega with default settings (https://www.ebi.ac.uk/Tools/msa/clustalo/). The WT sequence encoding the plastid targeting sequence is marked in green and the designed single-guide RNA (sgRNA) sequence and protospacer adjacent motif (PAM) in red. The position of genotyping primers is highlighted as arrows. Mutations are boxed in pink and translation start and stop codons are boxed in blue and yellow, respectively.

**Fig. S3.**
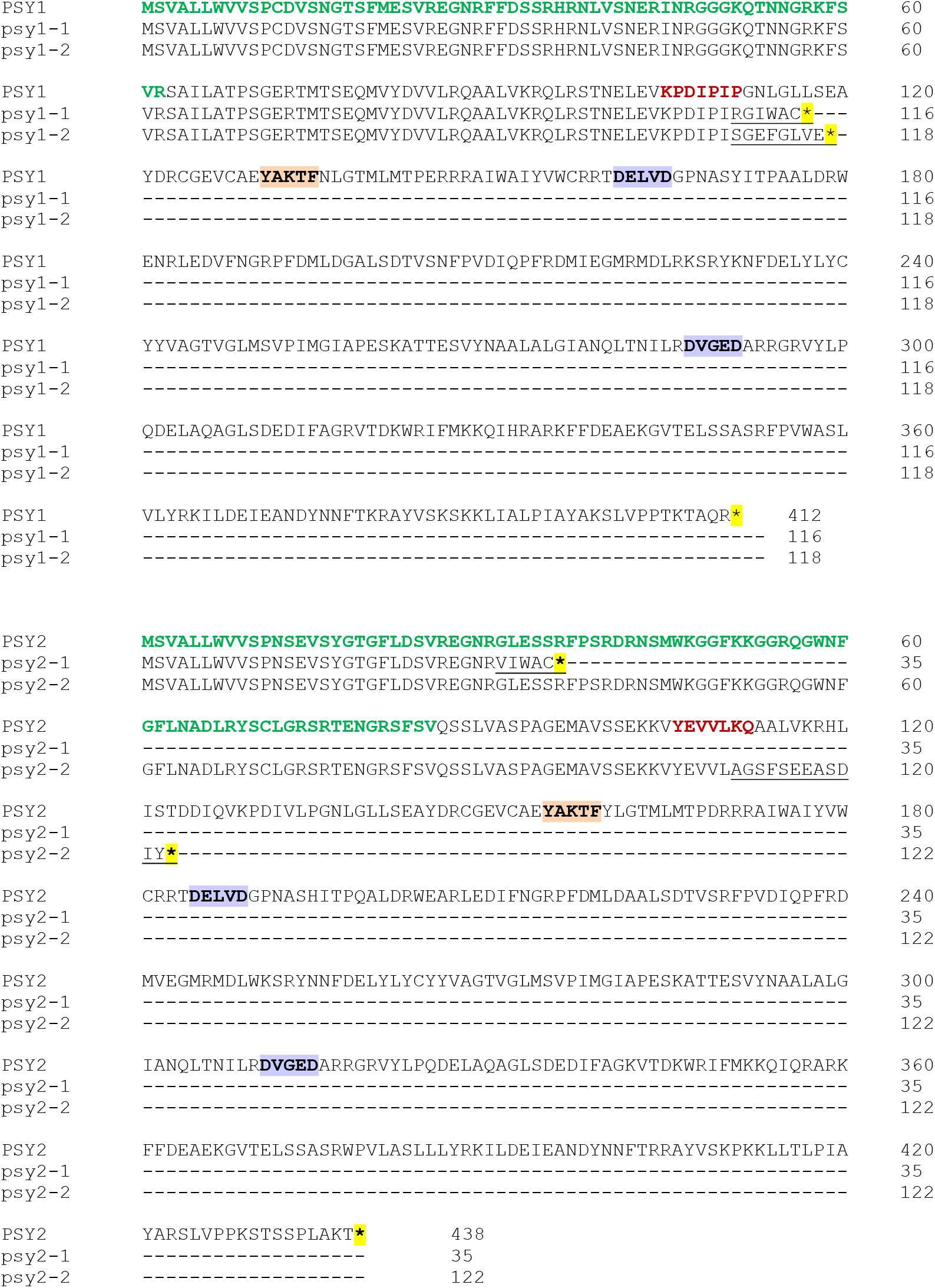
Protein sequence alignment of PSY1 and PSY2 sequences from WT and CRISPR mutants. Alignment was performed using Clustal Omega with default settings (https://www.ebi.ac.uk/Tools/msa/clustalo/). The WT sequences show the plastid targeting sequence marked in green and the position targeted by the designed single-guide RNA (sgRNA) sequence in red. They also show conserved domains pivotal to PSY function: hydrobofic flap (boxed in orange) and Asp-rich domains (boxed in purple). The sequence present in the different alleles as a consequence of their respective mutations is underlined. Translation stop codons are boxed in yellow.

**Fig. S4.**
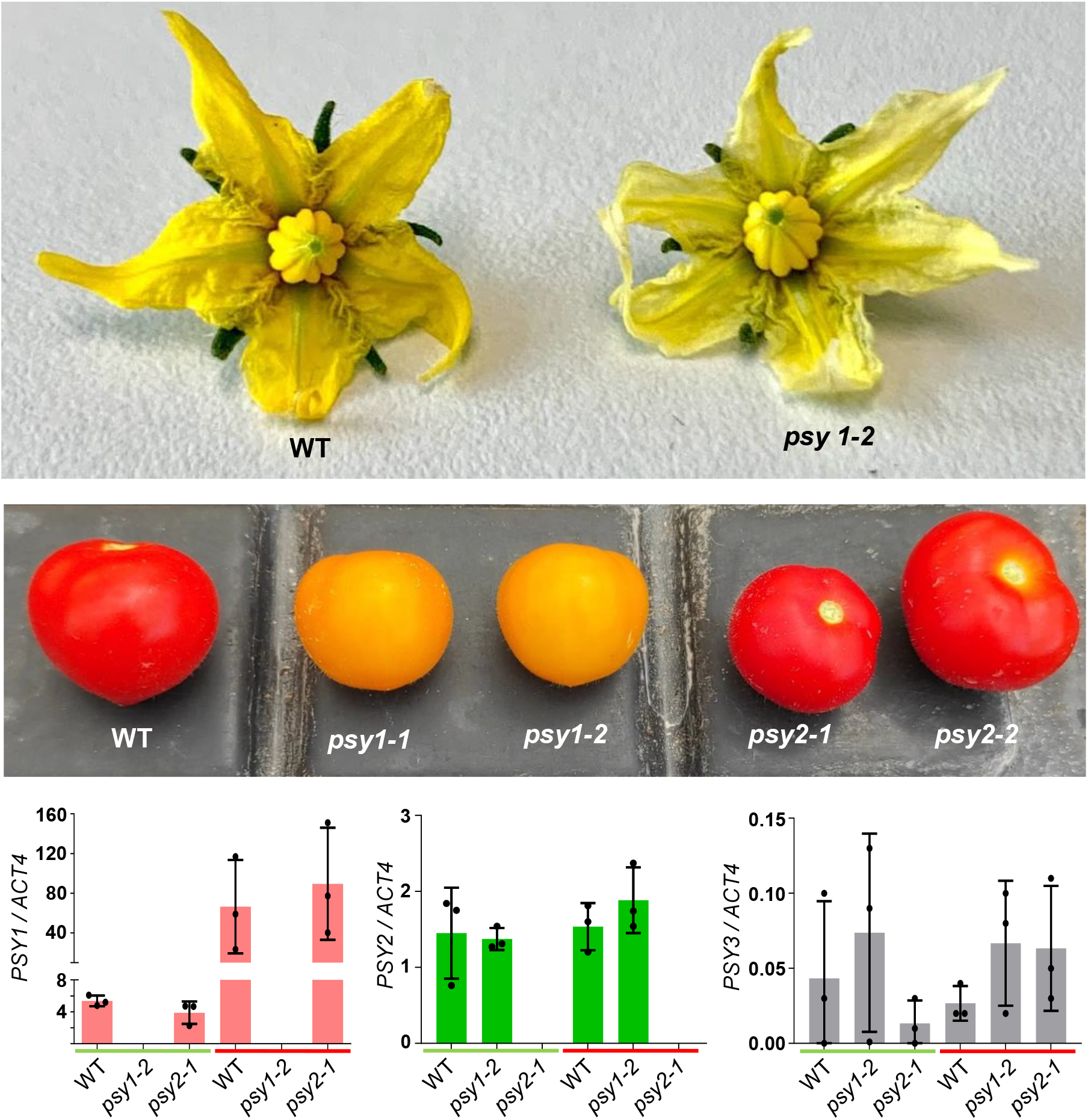
Representative phenotypes of tomato mutants defective in PSY1 or PSY2. The upper picture shows WT and *psy1-2* flowers in anthesis. The picture below shows representative ripe fruits from the indicated genotypes. The plots show the result of RT-qPCR analysis of *PSY1, PSY2* and *PSY3* transcript levels in pericarp tissue from WT and mutant fruit collected from the plant at MG (green bar) and ripe (B+6, red bar) stages. Individual values after normalization with the *ACT4* gene are shown together with the mean and SD of n=3 independent biological replicates.

**Figure S5.**
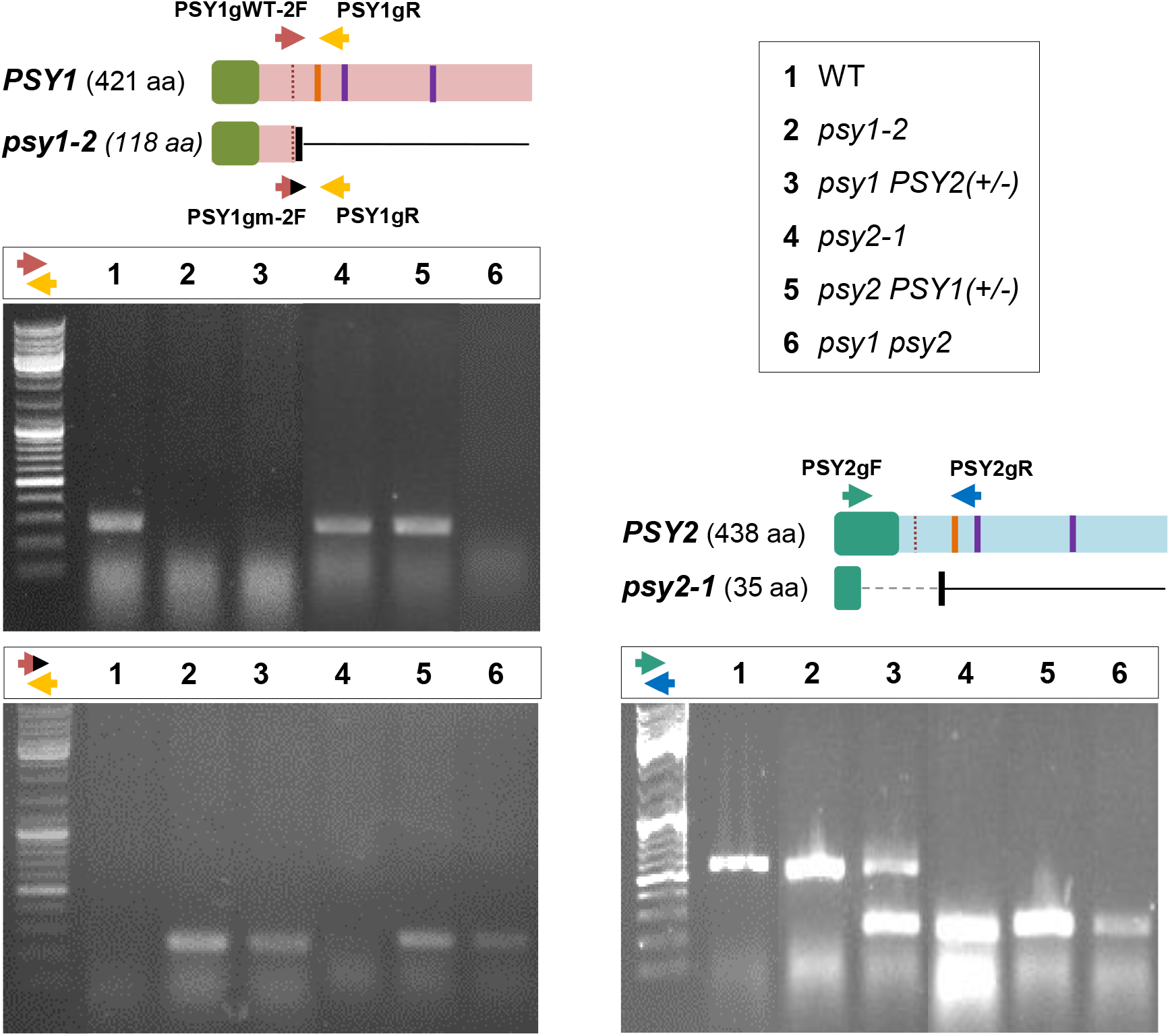
PCR genotyping of mutant alleles. The schemes representing the WT PSY1 and PSY2 proteins and the mutant versions generated in this work are described in Fig. 1B. Arrows represent the position of primers for genotyping. Agarose gel analysis of the results for the indicated genotypes resulting from the cross of *psy1-2* and *psy2-1* plants are shown.

**Figure S6.**
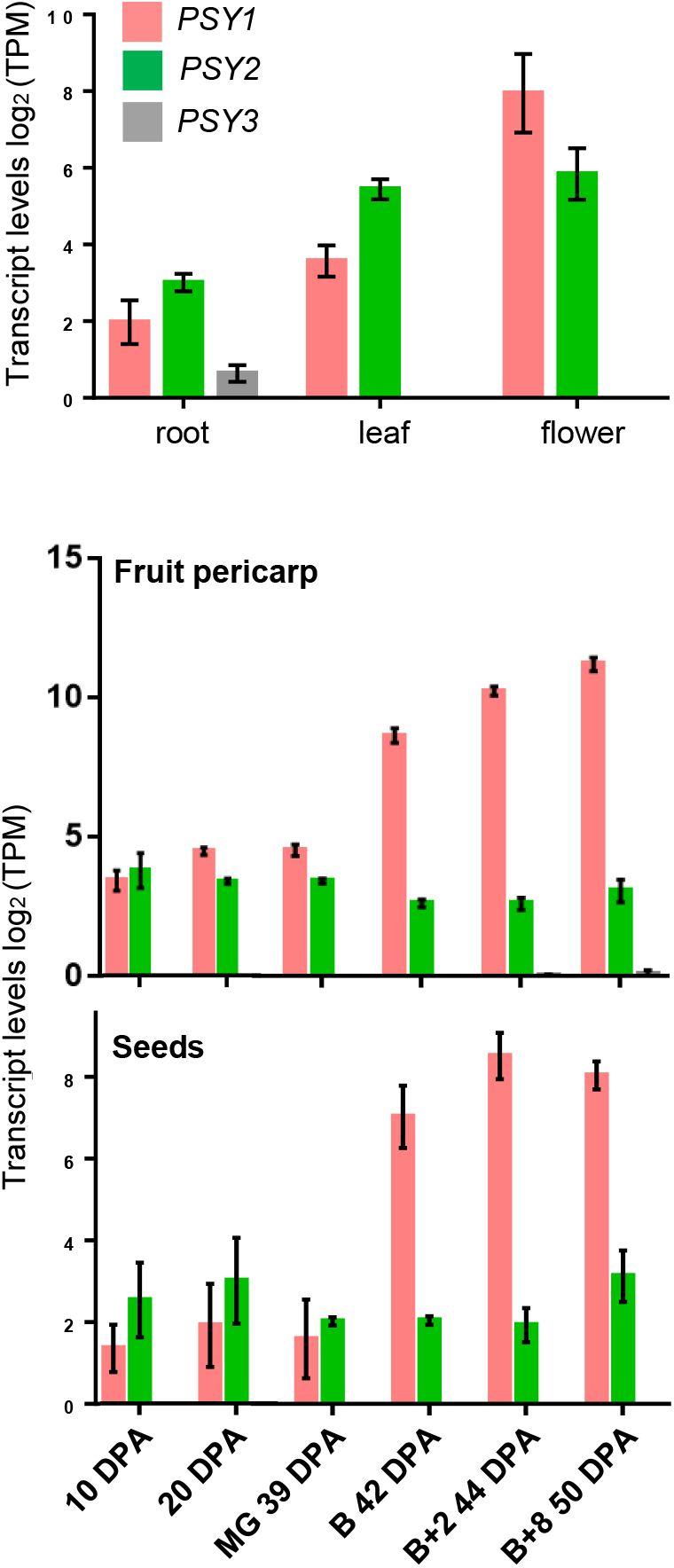
*PSY1, PSY2* and *PSY3* transcript levels in different tissues. Plots represent RNAseq data obtained from Genevestigator (https://genevestigator.com). Transcript levels are represented as log2 TPM (transcripts per million mapped reads). DPA, days post-anthesis; MG, mature green; B, breaker.

**Figure S7.**
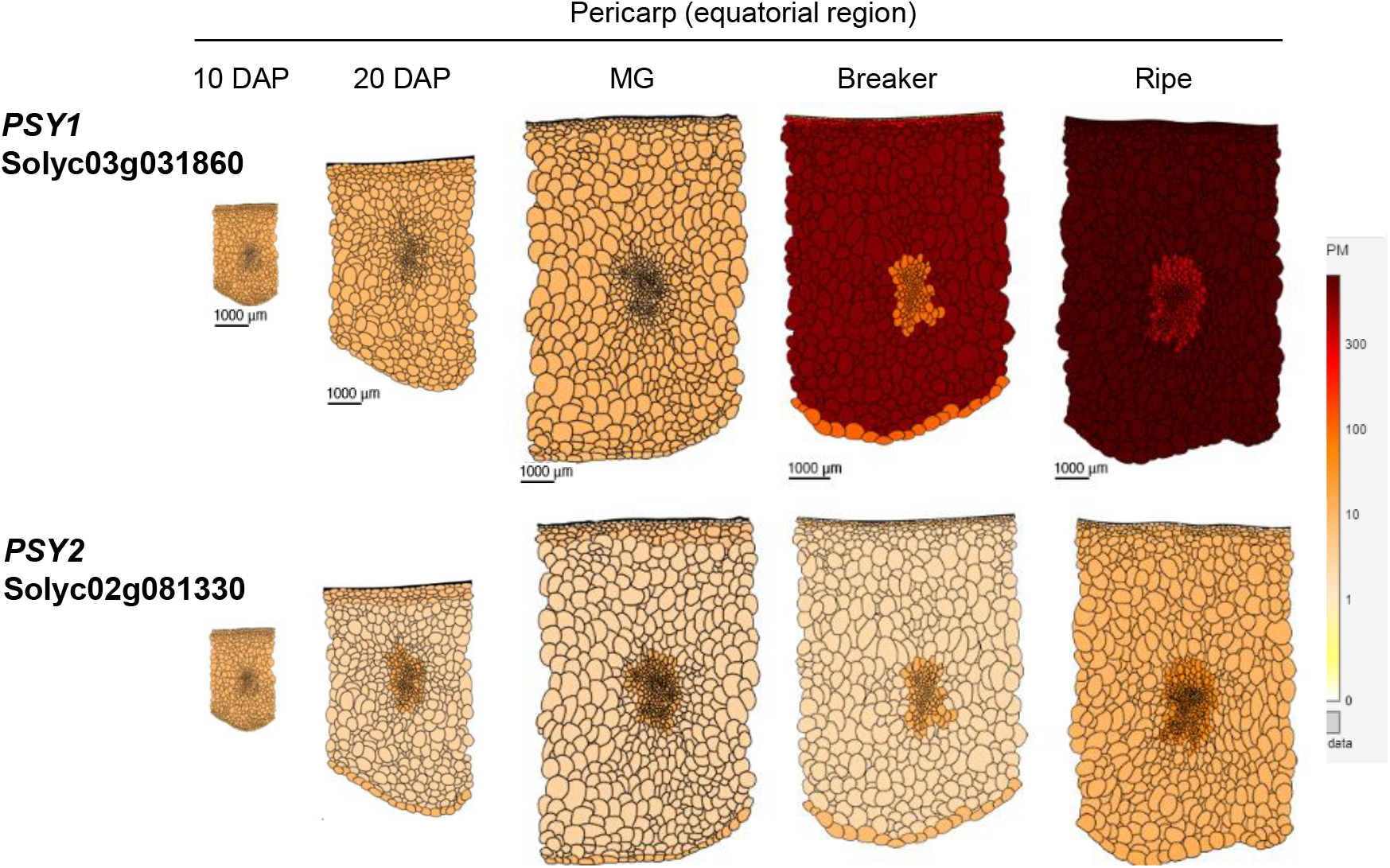
Expression profile of *PSY1* and *PSY2* in the fruit pericarp during development. Data were retrieved from the Tomato Expression Atlas’ expression viewer (https://tea.solgenomics.net/expression_viewer/input). DPA, days post-anthesis; MG, mature green.

**Figure S8.**
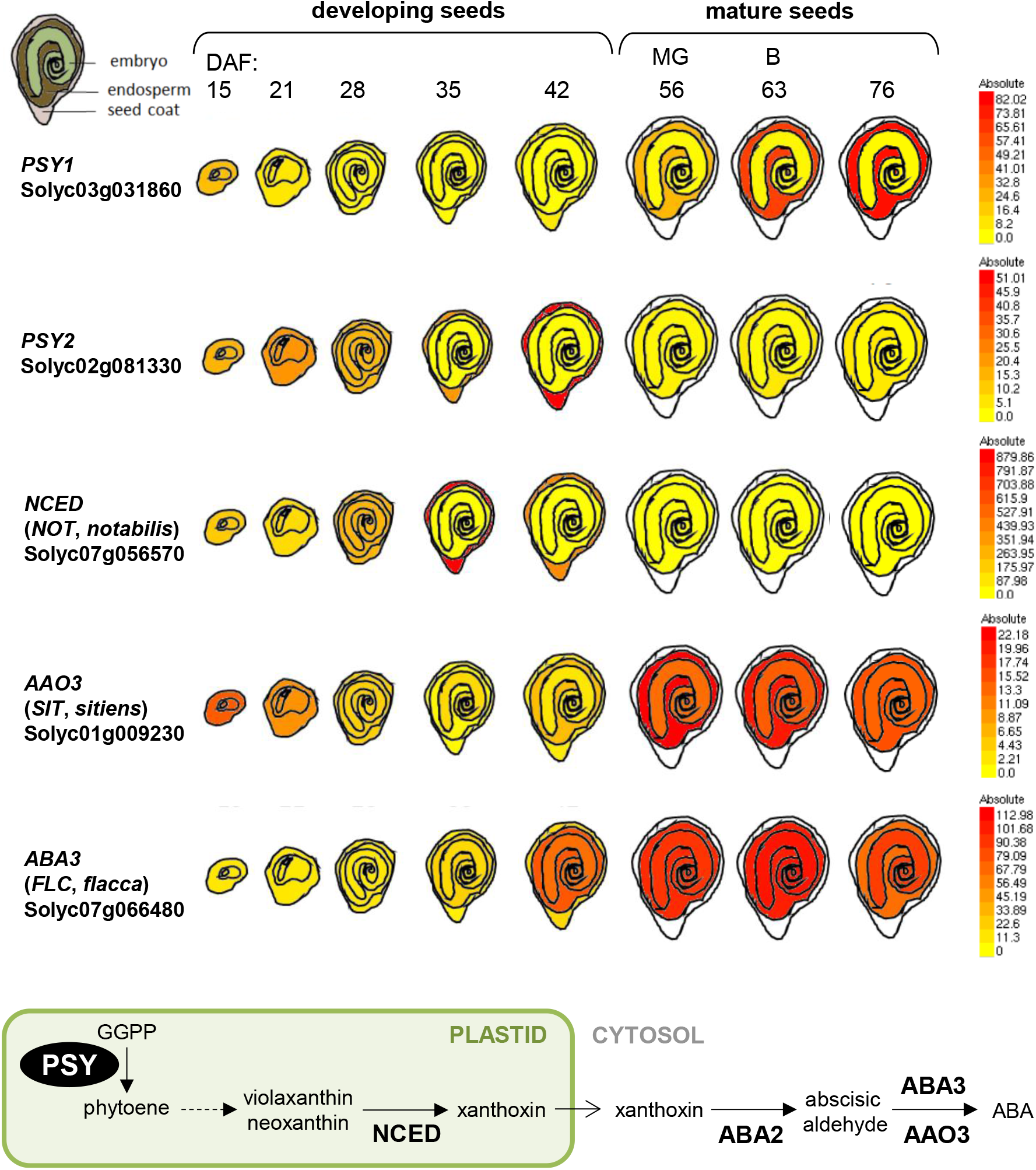
Expression profile of ABA biosynthetic genes in developing seeds. Data were retrieved from the Tomato eFP Browser (http://bar.utoronto.ca/efp_tomato/cgi-bin/efpWeb.cgi). DAF, days after flowering; MG, mature green; B, breaker. A schematic ABA biosynthesis pathway is also shown.

## Supplemental Tables

**Table S1.**
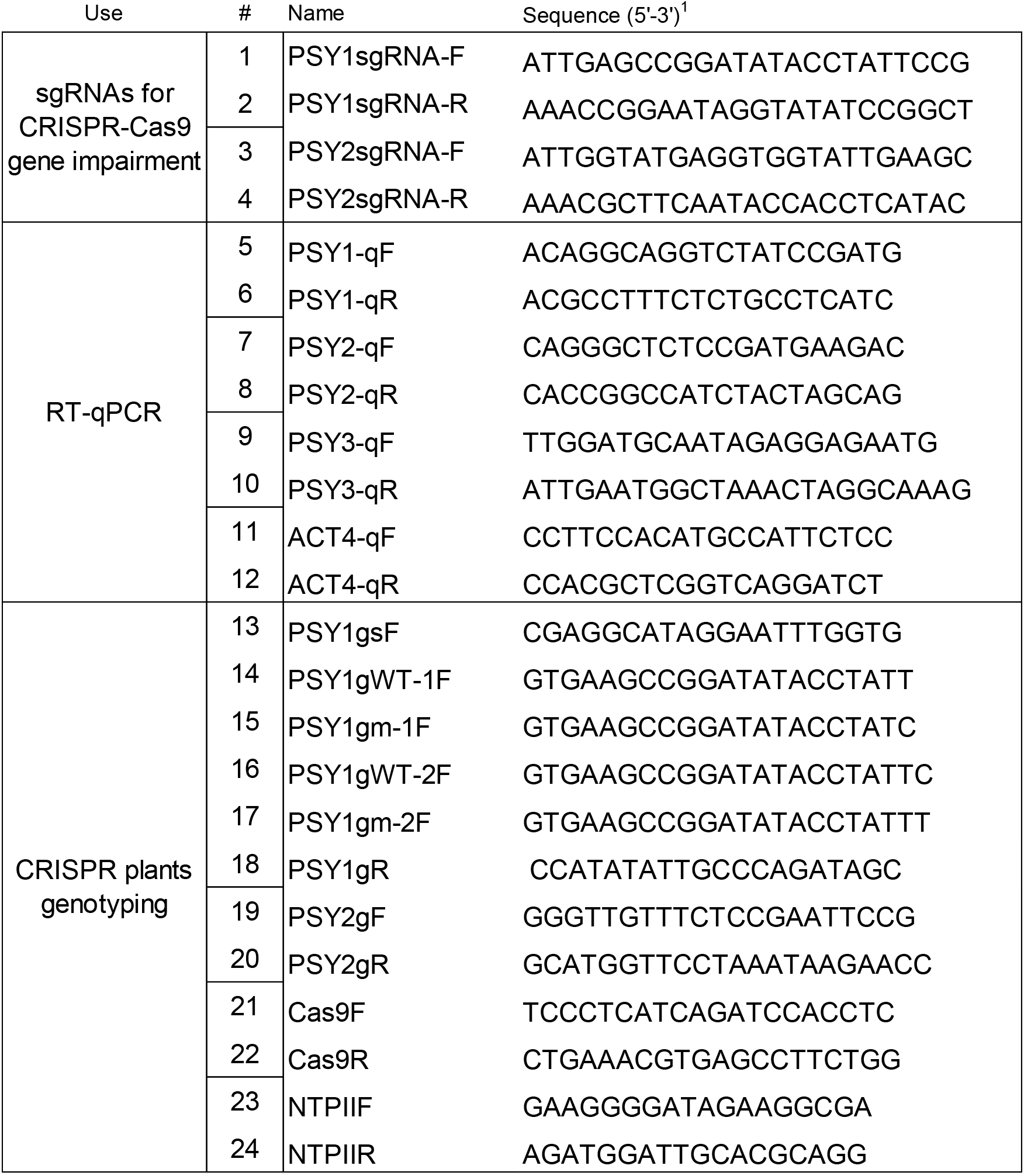
Primers used in this work.

**Table S2.** CRISPR-Cas9 constructs and cloning details.

| **Construct** | **Template** | **Primers** | **Sequence**<br>**cloned\*** | **Cloning method** | **Entry plasmid** | **Destiny**<br>**plasmid** |
| --- | --- | --- | --- | --- | --- | --- |
| *pEN-PSY1(sg1)* | - | 1 + 2 | PSY1<sub>1849-1868</sub> | *Bbs*I / T4 ligase | pENC1.1 | - |
| *pEN-PSY2(sg2)* | - | 3 + 4 | PSY2<sub>806-825</sub> | *Bbs*I / T4 ligase | pENC1.1 | - |
| *pDE-PSY1,2*<br>*(sg1+sg2)* | *pEN-PSY1(sg1)* +<br>*pEN-PSY2(sg2)* | - | PSY1<sub>1849-1868</sub> +<br>PSY2<sub>806-825</sub> | PSY1(sg1) *Mlu*I +<br>*Bsu*36I / T4 ligase<br>PSY2(sg2) Gateway | *pEN-PSY1(sg1)* +<br>*pEN-PSY2(sg2)* | pDE-Cas9 |

\* Numbers indicate the first and last nucleotide positions in the genomic DNA (see Fig. S1 for PSY1 and Fig. S2 for PSY2)

